# Spontaneous slow waves index multiple dimensions of recovery in Disorders of Consciousness

**DOI:** 10.64898/2026.09.30.755786

**Authors:** Alessia Ruyant Belabbas, Augustin Boullet, Dragana Manasova, Arthur Le Coz, Mélanie Valente, François-Xavier Lejeune, Jacobo Diego Sitt, Lionel Naccache, Benjamin Rohaut, Thomas Andrillon

## Abstract

Whether and how residual awareness is expressed in patients with Disorders of Consciousness (DoC) depends on the severity of brain damage and the transient functional state of the cortex at assessment. Classically, DoC are associated with electroencephalographic (EEG) slowing, marked by high-amplitude slow waves, even during the day. Yet, slow waves are also a hallmark of healthy sleep, where they are thought to serve restorative functions and to help maintain network homeostasis. Here, we asked whether slow waves in DoC can provide insights into the severity of cortical dysfunction or, conversely, into the preservation of residual awareness and recovery potential, by characterizing their spatial, temporal and morphological properties.

We analyzed 256-channel EEG recorded during the ‘Local-Global’ auditory paradigm in 236 DoC patients, including 123 in Unresponsive Wakefulness Syndrome (UWS), 62 in a Minimally Conscious State minus (MCS-) and 29 in MCS plus (MCS+). High-amplitude slow waves were detected as local events, and their density, amplitude and slopes were quantified scalp-wide and channel-wise, by mean level and temporal variability across the recording. Slow-wave features were related to concurrent (Coma Recovery Scale-Revised - CRS-R, ‘Local-Global’ paradigm) and follow-up (one-year Glasgow Outcome Scale-Extended - GOS-E) assessments of consciousness and functional recovery. Associations with MRI-derived measures of cortical integrity and structural connectivity, and differences between anoxic and traumatic etiologies, were also examined.

Slow waves were more frequent in less severe DoC patients (i.e., MCS) compared to UWS. Slow-wave slopes were also steeper frontally and increased at a faster rate in patients exhibiting higher levels of consciousness and/or markers of conscious access during the ‘Local-Global’ paradigm. More frequent and steeper slow waves at baseline further predicted recovery of consciousness and of functional independence at one year. At the individual level, slow-wave features classified states of consciousness above chance (AUC = 0.62 and 0.71 for MCS- and MCS+ versus UWS), whereas outcome prediction remained modest (0.56-0.58). These clinical associations had a structural basis: slow-wave occurrence was greater after traumatic than anoxic injury and scaled with the integrity of subcortico-cortical projection fibers and consciousness-related cortical networks.

Contrary to the classical view of delta activity as a non-specific marker of impaired consciousness, prominent spontaneous slow waves indexed behavioral and neurophysiological signs of awareness and better recovery. Beyond reflecting a propensity for pathological bistability, they may also signal the integrity of the networks required to generate them, together with their residual capacity for dynamic regulation.

**Highlights:**

1. Spontaneous slow waves were detected as discrete local events in 236 DoC patients, moving beyond time-averaged spectral measures of cortical slowing.
2. Slow-wave properties carried diagnostic information about the state of consciousness, whether defined behaviorally (CRS-R) or neurophysiologically (conscious access to auditory information in ‘Local-Global’ ERP paradigm).
3. Slow wave frequency and slope during the acute phase in the ICU predicted improvement of consciousness state and recovery of functional independence at one year.
4. Within-session ‘fatigue-like’ temporal modulation of slow waves increased with the level of consciousness recovery, being more pronounced in MCS+ than in UWS and MCS-patients.
5. Slow-wave expression scaled with subcortico-cortical fibers and cortical network integrity, and was greater after traumatic than anoxic injury, suggesting they signal preserved network integrity rather than damage alone.

## INTRODUCTION

Disorders of Consciousness (DoC) refer to a heterogeneous spectrum of clinical conditions characterized by varying degrees of impairment of awareness following severe acquired brain injury. They encompass a wide range of etiologies, including traumatic brain injury (TBI), anoxic damage, and stroke, and often present with complex clinical presentations involving multiple interacting pathophysiological mechanisms^1,2^. Accurately assessing a patient’s current level of consciousness (diagnosis) and predicting their potential for recovery (prognosis) remain major medical, scientific, and ethical challenges. Because conscious experience is inherently private and subjective^3^, its assessment in others necessarily relies on indirect evidence, most commonly verbal reports or purposeful behavior interpreted as signs of awareness^4^. This makes the evaluation of consciousness particularly difficult in non-communicating patients, whose repertoire of overt responses is often severely limited by motor, sensory, or cognitive impairments^4–7^. Distinguishing between the Unresponsive Wakefulness Syndrome (UWS), defined by wakefulness without behavioral evidence of awareness, and the Minimally Conscious State (MCS), in which patients may exhibit reproducible but inconsistent signs of conscious interaction with their environment, is particularly challenging, yet carries profound implications for prognosis, clinical management, and goals of care decisions^8–11^.

To reduce the risk of misdiagnosis and underestimation of patients’ level of consciousness, current international guidelines therefore advocate a multimodal approach^11,12^ combining behavioral assessments with structural and functional neuroimaging and neurophysiological techniques (e.g., MRI, PET, and EEG). These strategies have considerably improved diagnostic accuracy and prognostic prediction by capturing complementary dimensions of brain injury and residual brain function (e.g., structural integrity, cerebral metabolism, and large-scale brain network function)^13–15^.

One dimension, however, remains overlooked within this multidimensional framework: transient changes in functional brain state and cortical dynamics. DoC patients frequently exhibit marked fluctuations in arousal and behavioral responsiveness, both within and across days^16–20^. These fluctuations are widely recognized clinically^21^: the gold-standard Coma Recovery Scale-Revised^22^ (CRS-R) includes a standardized arousal facilitation protocol, repeated whenever arousal or responsiveness declines during assessment, and current guidelines recommend repeated behavioral evaluations across sessions to avoid missing behavioral signs of consciousness^11,12,23^. Fluctuations in vigilance or attention may also carry direct clinical consequences: beyond affecting behavioral diagnostic accuracy^23,24^, they may reduce the expression of residual cognitive processing during neurophysiological assessment, plausibly contributing to the limited sensitivity of active paradigms such as the ‘Local-Global’ task^25–27^. At the neurophysiological level, these fluctuations have been explored through the dynamics of patterns of functional connectivity obtained with fMRI^28^, or micro-states derived from EEG^29–31^. Together, these observations point to the clinical relevance of understanding the physiological mechanisms underlying these fluctuations, which are likely driven by transient changes in cortical dynamics.

In healthy individuals, such fluctuations in vigilance and responsiveness to external stimulation have been closely linked to the phenomenon of cortical slowing reflected by the intrusion of EEG slow waves into ongoing brain activity^32,33^. Although typically considered a hallmark of physiological sleep, slow waves can locally intrude into wakefulness as local sleep events and have been associated with fluctuations in arousal, attentional lapses, altered subjective experience (e.g., mind-wandering and mind-blanking), and reduced responsiveness to external stimuli^32,34–39^ (see ^33,40^ for reviews). Slow-wave occurrence is tightly linked to sleep homeostasis, increasing with sleep pressure^34–36,41^, and is thought to reflect transient neuronal off-periods reflecting local alterations in cortical dynamics and increased cortical bistability^34,37,42^.

Beyond this role in physiological state regulation, slow-wave activity can also directly signal structural brain injury. Pathological slow-wave activity has long been recognized as a common electrophysiological signature of brain injury^43–45^. More recently, focal stroke studies have shown that similar high-amplitude delta oscillations can emerge not only peri-lesionally but also within structurally intact, functionally connected cortex^46,47^, through diaschisis^48–50^. Experimental models of cortical deafferentation, including isolated cortical slabs^51^ and hemispherotomy^52^ have demonstrated that structural disconnection alone is sufficient to generate sleep-like slow waves during wakefulness in the disconnected cortex. Together, these findings indicate that the emergence of pathological slow waves depends not only on the site of injury but more generally on the disruption of structural and functional network integration.

This framework is particularly relevant to DoC, which arise from severe acquired brain injury and are characterized by profound structural and functional brain alterations across multiple spatial scales^1,2,53^. Diffuse cortical slowing is one of the most consistent electrophysiological features of DoC^54^. Importantly, these slow waves observed in DoC resemble those observed in slow-wave sleep and general anesthesia^55^, potentially reflecting a continuum between pathological and normal slow waves in DoC patients. Because they reflect a change in the cortical dynamics that support information processing and exchange across brain networks^40^, these slow waves are thought to be not just a marker of altered states of consciousness but a mechanism involved in the loss of consciousness^56^. Indeed, recent theoretical work has proposed that transient occurrences of sleep-like slow waves during wakefulness in DoC may contribute to fluctuations in responsiveness and potentially shape the subjective experience of patients with residual consciousness^27^. This is consistent with the broader pattern of injury in DoC: across etiologies, focal and diffuse damage to cortical, subcortical and white matter structures, notably in thalamo-cortical systems^53^, often disrupts the structural substrate underlying this large-scale exchange. DoC are correspondingly associated with reduced connectivity^26,57–59^ and diminished complexity^26,60–63^, both of which scale with the behavioral level of consciousness and could be linked to cortical slowing. Sleep-like slow waves may thus provide a particularly informative neurophysiological readout of the altered cortical dynamics that emerge from these structural and functional disruptions.

Slow-wave activity thus sits at the intersection of two key pathophysiological dimensions of DoC: as a single scalp EEG phenomenon, it may simultaneously index sleep-related changes in functional brain state and injury-related changes in structural brain integrity, making it a particularly promising neurophysiological target for investigating DoC. Yet slow-wave activity remains partially characterized in DoC. Most EEG studies have approached cortical slowing through spectral measures of delta power (0.5-4Hz), showing that increased low-frequency activity is associated with lower levels of consciousness^26,64–66^ and poorer clinical outcomes^67–69^. More recently, TMS-EEG studies further highlighted the potential relevance of slow-wave dynamics by showing that cortical responses to transcranial magnetic stimulation in behaviorally awake UWS patients are characterized by TMS-evoked sleep-like slow waves associated with spectral OFF periods^70^. While this establishes an important conceptual link between DoC and sleep-like cortical dynamics, whether spontaneous sleep-like slow waves occurring in ongoing EEG activity carry clinically relevant information remains largely unknown. Critically, these spectral approaches provide only a global, time-averaged measure of cortical slowing and overlook the discrete, event-like nature of slow waves. Consequently, they cannot capture the intrinsic properties and temporal dynamics of individual slow-wave events, which may carry distinct physiological information. An event-based characterization of slow waves may overcome some of these limitations by preserving their spatial, temporal and morphological diversity.

Here, we investigated the clinical relevance of characterizing slow waves as discrete, local neurophysiological events in a large cohort of patients with Disorders of Consciousness. To address these complementary aspects, we examined their associations with behavioral and functional neurophysiological markers of the state of consciousness, characterized their dynamic component by assessing their moment-to-moment temporal variability, examined their associations with etiological context and MRI-derived measures of structural integrity, and evaluated their ability to predict long-term cognitive outcome. Finally, we asked whether the multidimensional characterization of sleep-like slow waves could be leveraged to improve diagnostic and prognostic classifiers at the individual patient level.

## METHODS

### Patients Inclusion and Ethics

Patients admitted to Pitié-Salpêtrière Hospital between 2014 and 2022 for expert assessment of consciousness state and neuroprognostication following severe acquired brain injury were included. The cohort encompassed a broad range of etiologies of Disorders of Consciousness, including, but not limited to, traumatic brain injury, anoxia, stroke, and metabolic disorders, as well as varying degrees of chronicity.

This study was ancillary to a patient cohort approved by the Ethics Committee of the French Society of Intensive Care Medicine - SRLF; Paris, France (NEURO-DoC/HAO-006/20130409 and M-NEURO-DoC/NCT04534777, https://clinicaltrials.gov/ct2/show/NCT04534777) in accordance with French regulations governing routine care research and conducted in compliance with the principles of the Declaration of Helsinki. Written information was delivered to and signed by a patient’s surrogate. Patients who recovered consciousness were given the opportunity to withdraw from the study. Inclusion criteria were (1) presence of a DoC requiring expert evaluation from the DoC-NeuroICU team, (2) evidence of brain injury confirmed by neuroimaging (CT or MRI) and (3) age between 18 and 80 years. Exclusion criteria were (1) deep sedation (e.g., refractory status epilepticus or elevated intracranial pressure); (2) pre-existing severe neurodegenerative disease (e.g., Alzheimer’s disease); and (3) pregnancy.

### Multimodal assessment in the ICU

During a standardized multimodal assessment lasting approximately one week, patients underwent repeated neurological examination and behavioral evaluations using the Coma Recovery Scale-Revised (CRS-R^22^), alongside neuroimaging (MRI, DTI, CT, PET scans) and neurophysiological recordings (high-density EEG at bedside). EEG recordings were systematically acquired both at rest and during the ‘Local-Global’ auditory paradigm described below^25^.

### Follow-up and Outcomes

Following this assessment phase in hospitalization, patients were longitudinally followed at 6 months, 1 year, and 2 years. Outcome data were collected through structured telephone interviews with the patient’s current physician or a close relative. Consciousness outcome was assessed by a phone-guided CRS-R evaluation^22,71^ allowing classification into coma, Unresponsive Wakefulness Syndrome (UWS), Minimally Conscious State (MCS-/MCS+), or emergence from MCS (EMCS). Functional recovery was assessed using the Glasgow Outcome Scale-Extended (GOS-E^72,73^).

For analyses, the CRS-R state obtained at follow-up was compared to the CRS-R assessment performed closest in time to the EEG recording used for analysis during hospitalization. Patient trajectories were categorized at each time point as: ‘improved’ (e.g., UWS at hospitalization and MCS-at follow-up), ‘stagnated’ (e.g., MCS+ at both time points), or ‘deceased’. For some analyses, ‘stagnated’ and ‘deceased’ patients were grouped under a ‘non-improved’ category. Functional outcome based on GOS-E (scores 1-8) was dichotomized into ‘unfavorable’ (GOS-E < 4) and ‘favorable’ (GOS-E ≥ 4), the latter corresponding to patients who regained functional independence sufficient to be left alone for up to 8 hours during the day. Complementary analyses compared patients with favorable outcomes (i.e., recovered consciousness and autonomy) to those with unfavorable but non-fatal outcomes (conscious with severe disability or persistent DoC).

Analyses focusing on the prognostic value of slow waves using the aforementioned outcome metrics were restricted to patients assessed within 3 months of DoC onset (N=179), excluding chronic cases (N=57). Main analyses included all deaths irrespective of whether they followed spontaneous disease progression or withdrawal of life-sustaining therapies (WLST). Complementary analyses reported in the Supplementary Figures separately examine deaths occurring with and without WLST to assess the potential impact of self-fulfilling prophecy bias.

### EEG recordings: ‘Local-Global’ paradigm

As part of the multimodal assessment, patients’ brain activity was recorded using high-density EEG while they underwent the ‘Local-Global’ paradigm^25^, for approximately 30 minutes (8 blocks of 3-4 minutes interleaved with rest periods). This auditory oddball paradigm informs DoC patient diagnosis by relying on event-related potentials (ERPs) to assess the brain’s ability to detect and process auditory patterns at two hierarchical levels^74,75^. Neural responses to ‘local’ violations in the auditory sequence (‘local effect’) are thought to reflect low-level auditory processing involving non-conscious iconic memory and are typically associated with the mismatch negativity (MMN) and P3a ERP components. In contrast, responses to global violations (‘global effect’) are known to index higher-order processing involving working memory, and are associated with the P3b component, commonly interpreted as a neural marker of conscious access to auditory information. The presence of ERP effects was determined using established criteria^25^ allowing patient classification into three categories: ‘no effect’ (absence of both local and global effects), ‘local only’ (presence of a local effect without global effect), and ‘local + global’ (presence of both effects), the latter being interpreted as evidence of preserved conscious processing. Behavioral assessment using the CRS-R was performed concurrently with EEG acquisition to determine the clinical state of consciousness at the time of recording. High-density scalp EEG was recorded using 256-channel HydroCel Geodesic Sensor Net systems and a Net Amps 300 amplifier (Electrical Geodesics, Inc.), digitized at 250 Hz and referenced to the vertex. To ensure data quality, impedances were kept below 100kΩ.

### EEG Dataset Filtering and Preprocessing

The initial database comprised 426 EEG recordings acquired in DoC patients undergoing the ‘Local-Global’ paradigm. Recordings were excluded from further analyses if associated clinical metadata were missing (e.g., absence of a corresponding CRS-R), if they were acquired during transcranial direct current stimulation (tDCS) sessions or under sedation, or if technical issues were identified (e.g., missing ‘Local-Global’ events or incomplete or irregular paradigm structure). After exclusion of 107 recordings, 319 EEG files were retained for preprocessing (see flowchart, Fig. 1).

**Figure 1:**
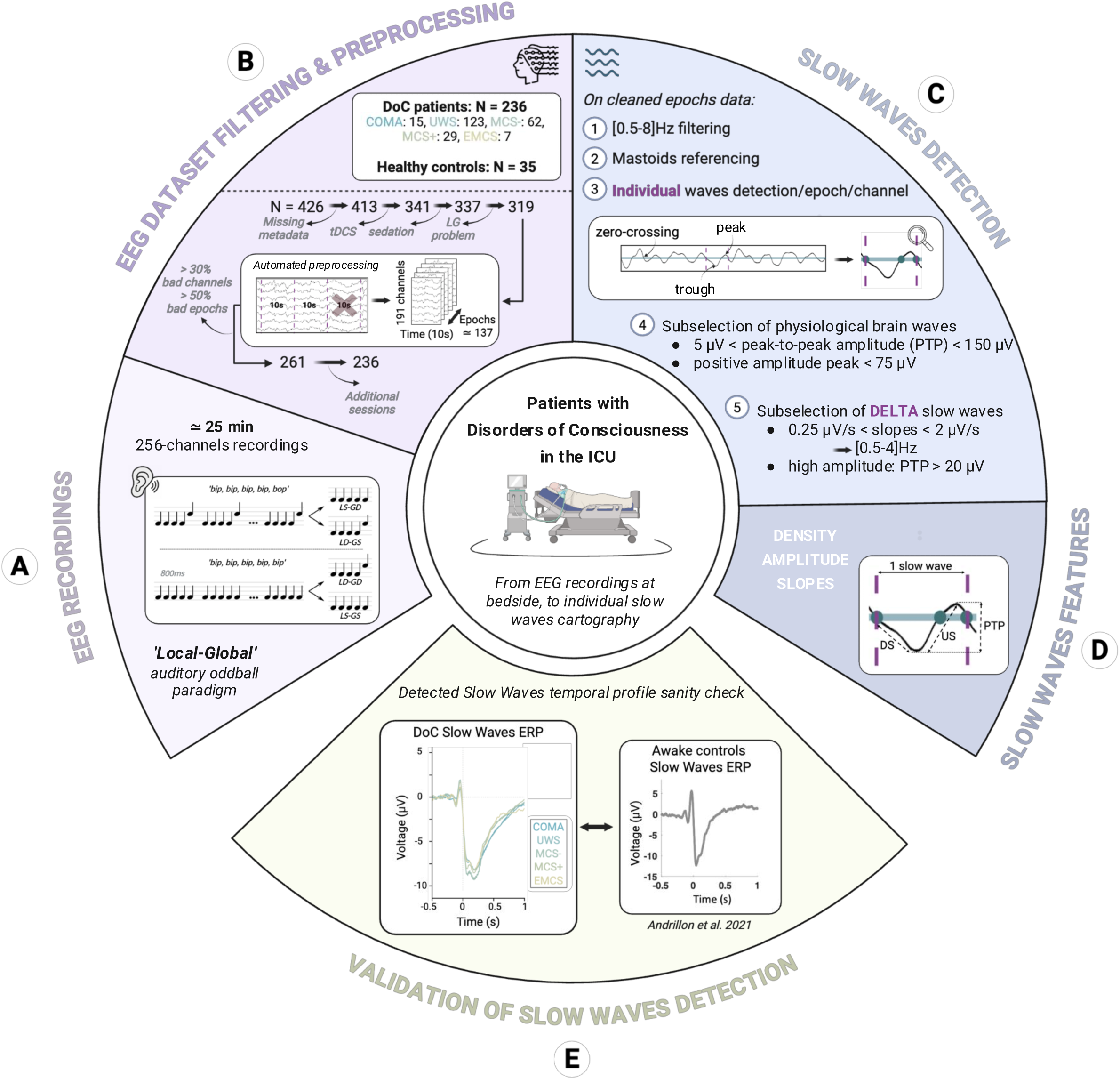
Experimental procedure. From high-density EEG recordings to individual slow waves exploration in DoC patients. **(A)** 256-channel EEG recordings dataset acquired at bedside in DoC patients undergoing the auditory oddball ‘Local-Global’ paradigm^25^. This paradigm, used for functional neurophysiological assessment, informs DoC diagnosis by probing the brain’s ability to detect auditory irregularities through ERPs. A ‘global effect’ (GD vs. GS; P3b) is typically interpreted as evidence of conscious integration of auditory information, relying on working memory, whereas a ‘local effect’ (LD vs LS; MMN and P3a) may testify to only low-level, non-conscious, auditory processing, relying on iconic memory. Illustration adapted from^141^ **(B)** EEG dataset subselection flowchart, including a data cleaning step using an automated preprocessing pipeline applied to continuous 10-s epochs (see Methods). **(C)** Overview of the algorithm used to detect high-amplitude individual delta slow waves. **(D)** Extracted features include slow-wave density (number of individual slow waves detected per channel, per 10-s epoch; in n/min), their peak-to-peak amplitude (PTP; in µV), and upward/downward slopes (US/DS, in µV/s) (see Methods). These metrics are subsequently averaged in the temporal dimension; per block of the ‘Local-Global’ task or across the entire recording (8 blocks); and in the spatial dimension; per channel or across all scalp channels. **(E)** Identified slow-wave temporal profile around Cz. Slow waves detected in DoC patients resemble the temporal profile of local sleep-like slow waves described in healthy individuals at wake^32^. The algorithm is well suited to detect DoC patients’ high-amplitude individual delta slow waves. DS = Downward slope; EMCS = Emergence from Minimally Conscious State; ERPs = Event-Related Potentials; GD = Global Deviant; GS = Global Standard; ICU = Intensive Care Unit; LD = Local Deviant; LG = ‘Local-Global’; LS = Local Standard; MCS = Minimally Conscious State; PTP = Peak-to-Peak amplitude; US = Upward slope; UWS = Unresponsive Wakefulness Syndrome. This figure was created with BioRender.

Preprocessing was performed using a standardized pipeline implemented in MNE-Python^76^. Electrodes located on the neck, cheeks and forehead were removed from the montage, leaving 191 channels for analysis. EEG signals were notch-filtered at 50 and 100 Hz (zero-phase FIR notch filter with a 1Hz transition bandwidth) to remove power line noise and band-pass filtered between 0.1 and 45 Hz using Butterworth IIR filters (6th-order high-pass, 8th-order low-pass). Continuous data were then segmented into non-overlapping 10-second epochs.

At the epoch level, baseline correction was applied using the entire 10-second epoch duration. An automated preprocessing pipeline based on the NICE toolbox^77^ was used for artifact detection and data cleaning. Channels were marked as bad if their signal amplitude exceeded 200 μV in more than 50% of epochs, while epochs were rejected if more than 10% of channels exceeded this voltage threshold. In addition, variance-based outlier detection was applied at both the channel and epoch levels: remaining electrodes and epochs with a variance Z-score greater than 4 were respectively marked as ‘bad’ and excluded, using an iterative procedure repeated four times. The remaining data were then re-referenced to the average.

Independent component analysis was performed on the retained channels (15 components; Infomax algorithm^78^, extended mode, random state = 97). Artifactual components (e.g., eye blinks, cardiac activity, channel noise) were identified using ICLabel^79^ and removed. Bad channels were subsequently interpolated from neighboring electrodes. Cleaned epochs were reassigned to their corresponding Local-Global block, ensuring that only ‘on task’ data were retained and inter-block intervals excluded from analyses.

Recordings with more than 30% bad channels and/or more than 50% rejected epochs were excluded, resulting in the removal of 58 additional files from the dataset. Finally, to ensure independence of observations, only one recording per patient was retained. When several recordings were available for a given patient, we kept the earliest one that passed quality control, thereby favoring assessments performed closer to DoC onset. This step led to the exclusion of 25 additional recordings.

The final dataset therefore comprised 236 EEG recordings from distinct DoC patients, with a median of 138 10-s epochs retained per recording [Q1-Q3: 113-163], corresponding to 23.0 minutes of data [Q1-Q3: 18.8-27.2; range: 12.5-38.8].

Clinical classification based on CRS-R collected at the time of recording included 15 coma, 123 UWS, 62 MCS-, 29 MCS+, and 7 EMCS patients. The cohort included 179 acute and 57 chronic patients (DoC onset > 3 months), with 73 post-anoxic (31%, cardiac arrest, hypoxia, low perfusion) and 40 post-traumatic (17%, TBI) etiologies, representing the two most prevalent causes of DoC in the cohort. In addition, 36 healthy controls underwent the same ‘Local-Global’ EEG protocol. One recording did not meet quality criteria and was excluded, resulting in 35 control recordings included in the analyses.

### Slow Waves Analyses

#### Slow Waves Detection

The detection of individual delta slow oscillations in DoC patients’ and healthy controls’ ‘Local-Global’ recordings used an approach initially devised to detect sleep slow waves in normal EEG recordings and later adapted to the detection of sleep-like slow waves in wakefulness^32,36,80^.

EEG signals from cleaned epochs data were re-referenced to the average of the two mastoids (E94 and E190) and band-pass filtered between 0.5 and 8Hz (delta-theta range) using a zero-phase finite impulse response filter with automatically determined length and transition bandwidths. The EEG filtered signal was then down-sampled to 100Hz (1000 samples per 10s epoch).

To identify individual waves, negative-to-positive and positive-to-negative zero crossings were detected at each channel and for each epoch, defined as time points at which the signal crossed the zero-voltage x-axis. Extrema between consecutive zero crossings (peak and trough of the wave) were located using the first derivative of the signal subsequently smoothed using a 5-sample moving average to improve robustness. Changes in signal direction (wave slope) were detected when the derivative exceeded a predefined slope threshold (0.1 μV/s). Troughs and peaks were identified at derivative sign changes (negative-to-positive and positive-to-negative, respectively) and retained only if their amplitudes were below and above 0 μV, respectively. Individual waves were thus defined as segments bounded by two consecutive negative zero-crossings, following a characteristic sequence: negative zero crossing (onset of the wave) → trough (maximum negative amplitude) → positive zero crossing → peak (maximum positive amplitude) → subsequent negative zero crossing (end of the wave) (see Fig. 1E).

For each candidate wave, the peak-to-peak amplitude (PTP) was computed as the voltage difference between the trough and the subsequent peak. To restrict detection to physiological brain events, only waves with a PTP between 5 and 150 μV and a positive peak amplitude below 75 μV were retained. Delta slow waves were then selected based on slope and amplitude criteria. Only waves with both downward and upward slopes between 0.25 and 2 μV/s (corresponding to 0.5-4Hz activity) were kept; with downward slope defined as the voltage decrease from the first negative zero crossing to the trough divided by the corresponding time interval and the upward slope as the voltage increase from the trough to the subsequent positive zero crossing divided by its duration. In addition, waves with a PTP below 20 μV were excluded, focusing analyses on the highest-amplitude delta slow waves.

#### Slow Waves Features

At each channel, and for every epoch within each block, the following features were extracted: slow-wave density (number of individual slow waves detected within the 10-s window and scaled by a factor of 6 to express values in min⁻¹), mean peak-to-peak amplitude (PTP; µV), and mean upward and downward slopes (US and DS; µV/s). These metrics were subsequently averaged across both temporal and spatial dimensions. Temporal averaging was performed at two levels: (i) within each block of the ‘Local-Global’ task (i.e., averaged across all epochs within a given block), and (ii) across the entire recording (i.e., averaged across all eight blocks). Spatial averaging was conducted either at the single-channel level for scalp topographies or across all scalp channels for global summary measures (e.g., boxplots).

#### Slow Waves Event Related Potential

Slow-wave temporal profile was obtained by locking and averaging EEG time series around detected slow waves. For all preprocessed 10-s epochs within each block and at each individual channel, data were re-epoched from -0.5 to +1s relative to the first negative zero-crossings, corresponding to the onset of the detected slow waves. Each wave-locked epoch was baseline-corrected using the -0.5 to -0.25s pre-onset interval. Resulting epochs were then averaged within each channel and block. For group-level visualization (Fig. 1E), slow-wave ERPs were averaged across channels of interest within each block (e.g., Cz), and subsequently averaged across all blocks from subjects within the same clinical state, as defined by the closest CRS-R assessment.

### MRI Analyses

#### MRI Acquisition and Preprocessing

MRI examinations were acquired as part of routine clinical multimodal imaging protocols using 1.5T and 3T scanners (General Electric HealthCare Technologies Inc., Chicago, USA and Siemens Healthineers, Erlangen, Germany). The imaging protocol included T1-weighted anatomical MRI (aMRI) acquired with a magnetization-prepared rapid gradient echo (MPRAGE) sequence, diffusion tensor imaging (DTI), and other conventional clinical sequences (axial T2-weighted, T2/FLAIR, and T2*-weighted imaging). Acquisition parameters, including repetition time, echo time, slice thickness, spatial resolution, and the number of diffusion-encoding directions, varied according to scanner model and clinical protocol. To account for this heterogeneity, all MRI data underwent standardized preprocessing and quality control procedures prior to feature extraction.

aMRI scans were preprocessed using FreeSurfer clinical software (v.7.4.1) with the recon-all-clinical implementation^81–83^. The automated pipeline included skull stripping, tissue segmentation, cortical surface reconstruction, and anatomical parcellation. Scans that failed automated segmentation or spatial normalization because of severe motion artefacts or extensive lesions were excluded from further analyses.

Quality control was also performed on all DTI images to ensure protocol compliance and exclude scans with significant motion or acquisition artefacts. Duplicate acquisitions from the same participant were also removed. DTI data were subsequently preprocessed using BrainQuant Software (www.braintale.eu; research version 2.0; Braintale, France), implementing preprocessing steps from the FMRIB Software Library (FSL), as previously described^84^. The preprocessing pipeline included correction for eddy-current-induced distortions and head motion, followed by brain extraction to remove non-brain tissue.

All aMRI and DTI scans were normalized to the Montreal Neurological Institute (MNI) standard space and further processed using automated pipelines, as described below.

#### Anatomical MRI: Cortical Thickness

Regional cortical thickness values were extracted using the FreeSurfer pipeline following cortical surface reconstruction and anatomical parcellation, as previously described^14^. Cortical measurements were subsequently organized according to the Schaefer-Yeo cortical atlas^85^ (100 regions of interest, 7 functional networks), comprising the Default Mode, Dorsal Attention, Salience/Ventral Attention, Somato-Motor, Visual, Limbic and Control (Frontoparietal) networks.

#### Diffusion Tensor Imaging: Fractional Anisotropy

Following the first preprocessing steps, a diffusion tensor model was fitted at each voxel to generate fractional anisotropy (FA) maps. FA maps were normalized to the MNI-152 template using linear and nonlinear transformations. Regional FA values were extracted for 21 predefined white matter tracts using the JHU-DTI-SS white matter atlas^86^, derived from the ICBM-DTI-81 atlas, and further grouped into projection, commissural, associative, and brainstem fibers categories. To account for inter-sequence variability arising from heterogeneous acquisition protocols, mean FA values were normalized by scaling each patient’s measurement to the mean FA of a normative control dataset acquired using identical imaging protocols.

### Statistics

Statistical analyses were conducted using linear models or linear mixed-effects models when repeated observations within subjects were included. Analyses were performed in R (version 4.4.2, R Core Team; *lm*, *lme4*, *lmerTest*, *emmeans* and *emtrends* packages), interfaced with Python via *rpy2*. For mixed-effects models, parameters were estimated using restricted maximum likelihood (REML), with Satterthwaite approximations used to estimate degrees of freedom. Model convergence was ensured using the *bobyqa* optimizer with extended iteration limits.

Fixed effects were modeled as categorical factors, except for Block in Fig. 3B, which was treated as a continuous numeric predictor to estimate linear trends over time, with block-wise changes normalized to Block 1.

Main effects of fixed factors and their interactions were assessed using omnibus analysis of variance (ANOVA) and are reported as F-values. Post hoc pairwise contrasts were computed on model-estimated marginal means (EMMs) and are reported as t-values. Resulting p-values were corrected for multiple comparisons using the Benjamini-Hochberg false discovery rate (FDR) procedure.

To assess the spatial distribution of effects across the scalp, the same models were fitted independently at each electrode, allowing the extraction of the corresponding channel-wise F-or t-values. Cluster-based permutation statistics implementing the Freedman-Lane procedure^87^ were then used to control for multiple comparisons across electrodes, while accounting for the spatial dependency structure of EEG data. Briefly, reduced models excluding the effect of interest were first fitted, and residuals were permuted across observations before reconstructing surrogate datasets and refitting the full model. For each of 1,000 permutations, the maximum cluster-level statistic (defined as the sum of absolute F- or t-values within each spatial cluster) was retained to build the null distribution. Spatial clusters were formed by identifying electrodes exceeding an uncorrected two-tailed p<0.05 cluster-forming threshold and grouping neighboring electrodes according to the MNE implementation of EEG sensor adjacency based on Delaunay triangulation. Cluster-level significance was then determined by comparing observed cluster-level statistics against the permutation-derived null distribution, controlling the family-wise error rate at α=0.05.

To examine the relationship between sleep-like slow-wave intrusion and the preservation of underlying brain anatomy (Fig. 5), correlations between MRI metrics (e.g., fractional anisotropy, cortical thickness) and slow-wave properties (e.g., density) were assessed using Spearman rank correlation coefficients (ρ), given the non-normal distribution of the variables and limited sample sizes. P-values were corrected for multiple comparisons using the FDR procedure. Differences in correlation strength between groups were assessed by transforming the correlation coefficients using Fisher’s z and comparing the resulting z-scores using the standard error for independent samples, with the Bonett-Wright variance correction applied to rank-based coefficients. In the figures, regression lines and their 95% confidence bands were derived from ordinary least squares fits to the raw data and are shown for visualization purposes only; all reported statistics are rank-based.

For patient-level diagnostic and prognostic classifiers, discriminant analyses were performed in R (version 4.4.2; R Core Team, 2023). Eight slow-wave metrics were extracted. For each metric, values were first averaged across channels within each recording block and then summarized across the eight recording blocks using either the mean or standard deviation, the latter capturing temporal variability. First, pairwise discriminant analyses between states of consciousness (MCS-, MCS+, and UWS) were performed separately for each metric. Thresholds were determined using the Youden index, and corresponding sensitivity and specificity were estimated with bootstrap confidence intervals based on 2,000 bootstrap replicates using the *pROC* R package (v1.18.5). A multivariate analysis combining all eight metrics was then performed using two-component partial least squares discriminant analysis (PLS-DA) with the *mixOmics* R package (v6.26.0)^88^. Metrics were ranked according to their discriminative contribution based on Variable Importance in Projection (VIP) scores and loading values. Model performance was assessed using ROC analysis. AUC values were obtained from the PLS-DA models using the *performance* function with 5-fold cross-validation repeated 20 times. Intervals around the AUC estimates were calculated as the mean AUC ±1.96 × the standard deviation across cross-validation repetitions. For visualization, the sensitivity and specificity of individual metrics were also displayed on the ROC curves. The same analysis was repeated for the prognostic outcomes: 1-year GOS-E (unfavorable vs favorable) and 1-year CRS-R evolution (improved vs not-improved) on the acute patients’ cohort (<3 months since DoC onset).

All statistical tests were two-sided, and statistical significance was set at p < 0.05 or at the corresponding multiple-comparison-corrected p < 0.05, as applicable.

## RESULTS

### Slow waves differentiate CRS-R subgroups and DoC etiologies

#### CRS-R subgroups

To assess the potential diagnostic value of slow waves in Disorders of Consciousness, we first investigated the relationship between slow-wave properties and CRS-R-defined states of consciousness (Fig. 2A).

**Figure 2:**
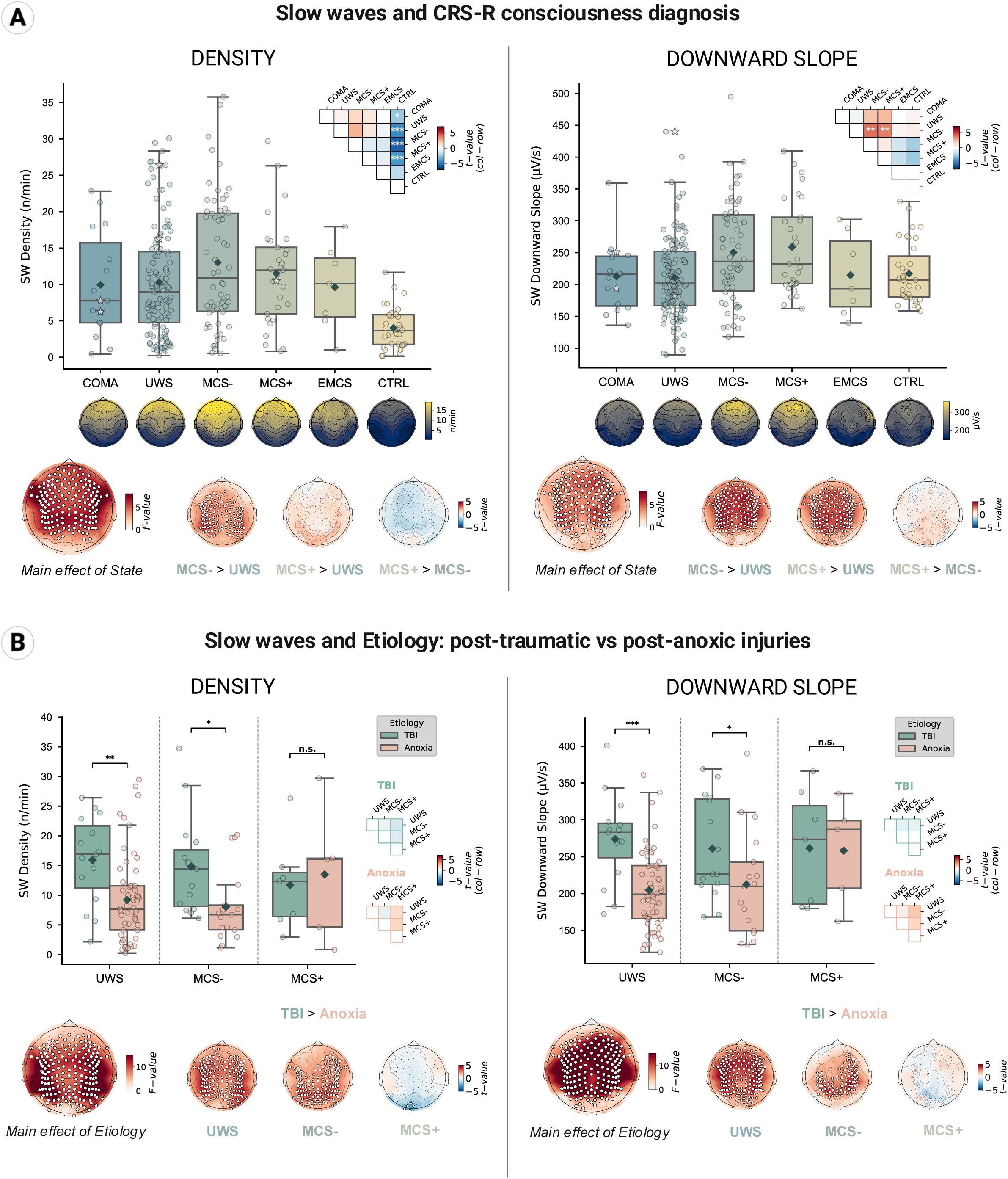
Slow-wave properties vary with clinically-defined consciousness state and etiology. Slow-wave metrics differ according to Coma Recovery Scale-Revised classification **(A)** and between post-traumatic and post-anoxic etiologies **(B)**. Distribution of slow wave Density (left) and Downward Slope (right), averaged across all channels and ‘Local-Global’ blocks, are shown for each consciousness state. Boxplots show the median (central line) and interquartile range (Q1-Q3, box edges), with whiskers extending to the most extreme values within 1.5 × IQR of the box edges. Group-level means are indicated by grey diamonds, while overlaid scatterplots display individual data points (circles). Boxplots are arranged according to increasing levels of consciousness recovery in DoC patients, from COMA to EMCS. CTRL refers to healthy controls. The triangle matrix displays pairwise comparisons between states, derived from linear models. Colors represent the t-statistics associated with each contrast (column - row), while asterisks indicate FDR-corrected significance levels (*: p < 0.05, **: p < 0.01, ***: p < 0.001). Asterisks above the boxplots indicate FDR-corrected between-etiology differences within each state, using the same significance thresholds; n.s., non-significant. Scalp topographies displayed above each boxplot illustrate the spatial distribution of the corresponding slow-wave metric. Bottom panels display statistical topographies of F-values (main effects) and t-values (pairwise contrasts). Channels showing statistically significant effects after CBPS correction (p < 0.05) are marked with white dots. CBPS = Cluster Based Permutation Statistics; CRS-R = Coma Recovery Scale-Revised; SW = Slow Waves; TBI = Traumatic Brain Injury.

Our analyses revealed that both scalp-averaged slow-wave density and downward slope differed across clinically defined consciousness states (linear model main effects: F_dens_ = 6.94, ^FDR^p_dens_ < 0.001; F_dslope_ = 4.75, ^FDR^p_dslope_ < 0.001). Pairwise contrasts showed that healthy controls exhibited lower slow-wave density than all DoC groups, except EMCS patients (t_dens(EMCS>CTRL)_ = 1.84, ^FDR^p_dens(EMCS>CTRL)_ = 0.17), consistent with the slowing of cortical dynamics commonly observed in altered states of consciousness. Slow-wave density tended to follow a non-linear, bell-shaped distribution across consciousness levels, with maximal values observed in intermediate stages of recovery. Accordingly, MCS- patients tended to display higher scalp-averaged density than UWS patients, although this difference did not survive correction for multiple comparisons (t_dens(MCS->UWS)_ = 2.37, ^FDR^p_dens(MCS->UWS)_ = 0.055, p_dens(MCS->UWS)_ = 0.019). Additionally, both MCS- and MCS+ patients displayed steeper slow-wave downward slopes than UWS patients (t_dslope(MCS->UWS)_ = 3.86, ^FDR^p_dslope(MCS->UWS)_ = 0.002; t_dslope(MCS+>UWS)_ = 3.54, ^FDR^p_dslope(MCS+>UWS)_ = 0.004), whereas no significant difference was observed between these two MCS subgroups (t_dslope(MCS+>MCS-)_ = 0.57, ^FDR^p_dslope(MCS+>MCS-)_ = 0.92).

Topographical analyses showed maximal slow-wave density over fronto-central regions (Fig. 2A), in line with reports that slow waves preferentially originate in frontal areas and propagate along an antero-posterior axis during physiological sleep^89,90^. Cluster-based permutation statistics (CBPS) identified diffuse effects of consciousness state on both density and downward slope, with strong main effects extending across large portions of the scalp (^CBPS^p_dens_ < 0.001, 176/191 electrodes, mass = 1010.87; ^CBPS^p_dslope_ < 0.001, 146 electrodes, mass = 613.03). Post hoc contrasts revealed widespread increases in high-amplitude slow-wave intrusions (^CBPS^p_dens(MCS->UWS)_ < 0.001, 120 electrodes, mass = 297.17) and downward slopes (^CBPS^p_dslope(MCS->UWS)_ < 0.001, 148 electrodes, mass = 507.48) in MCS- patients compared to UWS, as well as steeper downward slopes in MCS+ relative to UWS patients (^CBPS^p_dslope(MCS+>UWS)_ < 0.001, 152 electrodes, mass = 461.73). No significant cluster was found when contrasting MCS- and MCS+ patients, neither for density nor for downward slope.

#### DoC etiologies

We next investigated whether the relationship between spontaneous slow-wave intrusions and consciousness state depended on lesion context (Fig. 2B).

We observed a strong main effect of etiology on slow-wave density averaged across all channels (F_dens_ = 13.13, ^FDR^p_dens_ < 0.001), confirmed by topographical analyses with a cluster covering nearly the entire scalp (^CBPS^p_dens_ < 0.001, 177 electrodes, mass = 2084.60). Although the interaction with CRS-R consciousness state was not detected when averaging across the scalp (F_dens_ = 1.66, ^FDR^p_dens_ = 0.19), channel-wise analyses revealed a significant State × Etiology interaction over a posterior cluster (^CBPS^p_dens_ < 0.001, 27 electrodes, mass = 109.56) (Supplementary Fig. 1), indicating that the etiological effect was regionally modulated by the level of consciousness. Post hoc contrasts showed higher slow-wave density in post-traumatic relative to post-anoxic patients within both the UWS (t_dens(TBI>Anoxia)UWS_ = 3.11, ^FDR^p_dens (TBI>Anoxia)UWS_ = 0.002) and MCS- groups (t_dens(TBI>Anoxia)MCS-_ = 2.57, ^FDR^p_dens (TBI>Anoxia)MCS-_ = 0.011), whereas no effect was observed in MCS+ patients (t_dens(TBI>Anoxia)MCS+_ = -0.41, ^FDR^p_dens(TBI>Anoxia)MCS+_ = 0.69) scalp-wide. Topographically, this difference was diffuse in UWS patients (^CBPS^p_dens(TBI>Anoxia)UWS_ < 0.001, 165 electrodes, mass = 476.25) and remained widespread but spared left fronto-central regions in MCS- patients (^CBPS^p_dens(TBI>Anoxia)MCS-_ < 0.001, 143 electrodes, mass = 363.24). In MCS+ patients, only a small posterior cluster reached significance, with the opposite direction (higher density in post-anoxic patients; ^CBPS^p_dens(TBI>Anoxia)MCS+_ < 0.001, 5 electrodes, mass = 13.1), overlapping with the interaction cluster reported above.

Consistent with density findings, downward slope was also influenced by etiology (F_dslope_ = 17.33, ^FDR^p_dslope_ < 0.001), with steeper slopes in post-traumatic than post-anoxic patients in UWS (t_dslope(TBI>Anoxia)UWS_ = 3.89, ^FDR^p_dslope(TBI>Anoxia)UWS_ < 0.001) and MCS-(t_dslope(TBI>Anoxia)MCS-_ = 2.24, ^FDR^p_dslope(TBI>Anoxia)MCS-_ = 0.027), but not MCS+ groups (t_dslope(TBI>Anoxia)MCS+_ = 0.09, ^FDR^p_dslope(TBI>Anoxia)MCS+_ = 0.93). Topographically, the main effect extended across most of the scalp (^CBPS^p_dslope_ < 0.001, 166 electrodes, mass = 2709.32) and within-state contrasts confirmed a diffuse difference in UWS patients (^CBPS^p_dslope(TBI>Anoxia)UWS_ < 0.001, 166 electrodes, mass = 536.08), that was restricted to centro-parietal regions in MCS-patients (^CBPS^p_dslope (TBI>Anoxia)MCS-_ < 0.001, 91 electrodes, mass = 247.96). A significant State × etiology interaction emerged over a small left temporo-parietal cluster (^CBPS^p_dslope_ = 0.007, 4 electrodes, mass = 21.3) reflecting divergent patterns across etiologies: post-anoxic MCS+ patients showed steeper downward slopes than both UWS and MCS- patients over overlapping centro-parietal regions, consistent with the inverted U-shape profile (Fig. 2A), whereas post-traumatic MCS- patients showed shallower slopes than UWS patients over a small left fronto-temporal cluster, and more generally a tendency for gradual decrease in slow wave expression as consciousness level increases (Supplementary Fig. 1).

Overall, etiology exerted a robust influence on slow-wave expression, with post-traumatic patients exhibiting higher slow-wave density and steeper downward slopes than post-anoxic patients, particularly in lower levels of consciousness.

### Slow waves capture functional heterogeneities across and within DoC subgroups

To investigate whether slow waves reflect ongoing functional brain state, we compared slow-wave density and properties according to functional subgroups based on each individual’s ‘Local-Global’ ERP responses (Fig. 3A). These subgroups reflect neurophysiological evidence of auditory information processing ranging from low-level automatic novelty detection (MMN-like ‘local effect’) to conscious access to delivered stimuli (P3b-like ‘global effect’).

**Figure 3:**
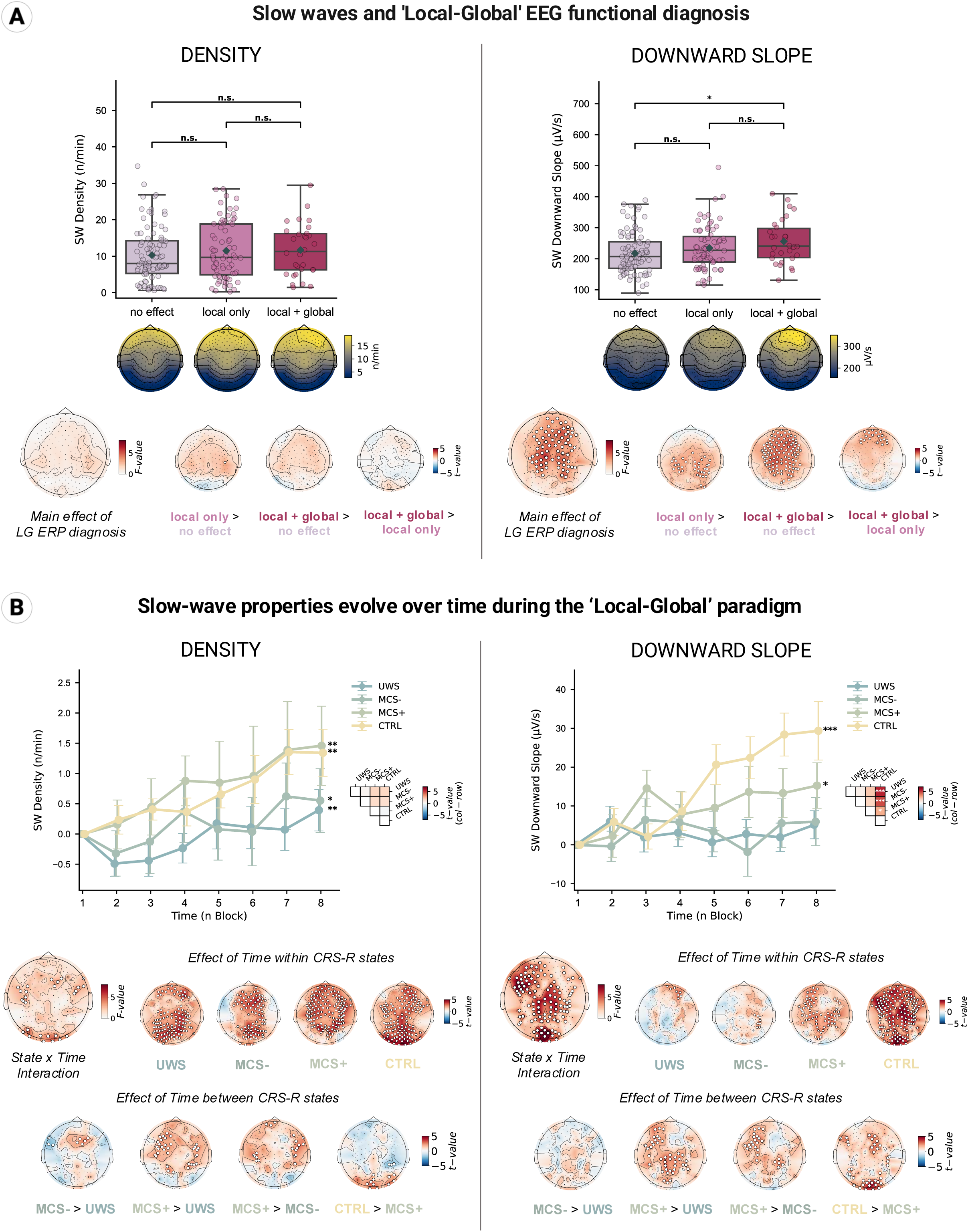
Slow-wave properties vary with neurophysiological functional state. Slow-wave metrics differ according to ERP evidence of conscious auditory processing during the ‘Local-Global’ paradigm **(A)**. Distributions of slow wave Density (left) and Downward Slope (right), averaged across all channels and ‘Local-Global’ blocks, are shown for each functional state. Boxplots are organized according to hierarchical levels of auditory processing: ‘no effect’, corresponding to the absence of both local and global ERP effects and indicating no evidence of auditory processing; ‘local only’, corresponding to the presence of a local ERP effect without a global effect, reflecting low-level non-conscious auditory processing; and ‘local + global’, indicating the presence of both ERP effects, testifying conscious integration of auditory information. Boxplots show the median (central line) and interquartile range (Q1-Q3, box edges), with whiskers extending to the most extreme values within 1.5 × IQR of the box edges. Group-level means are indicated by grey diamonds, while overlaid scatterplots display individual data points. Statistical annotations above the boxplots indicate FDR-corrected differences between ‘Local-Global’ functional states, derived from linear models (*: p < 0.05, **: p < 0.01, ***: p < 0.001, n.s.: non-significant). Scalp topographies displayed above each boxplot illustrate the spatial distribution of the corresponding slow wave metric. Bottom panels display statistical topographies of F-values (main effect) and t-values (pairwise contrast). **(B)** Slow-wave properties evolve over time during the ‘Local-Global’ paradigm. Slow wave metrics are shown across successive experimental blocks, probing temporal changes in slow waves activity during prolonged task exposure, within each clinical state. Line plots show slow wave Density (left) and Downward Slope (right), averaged across all channels and normalized to Block 1, across successive experimental blocks from Block 1 (start) to Block 8 (end). Group-level means are shown as connected lines, with error bars indicating standard error of the mean. Asterisks displayed to the right of the plots indicate FDR-corrected blocks effects (linear trend), derived from linear mixed-effects models (*: p < 0.05, **: p < 0.01, ***: p < 0.001). The triangle matrix displays pairwise comparisons between CRS-R states. Colors represent the t-statistics associated with each contrast (column - row), while asterisks indicate FDR-corrected significance levels using the aforementioned thresholds. Bottom panels display statistical topographies of F-values (interaction effect) and t-values (within state block effects and pairwise contrasts between states). **(A-B)** Channels showing statistically significant effects after CBPS correction (p < 0.05) are marked with white dots. CBPS = Cluster Based Permutation Statistics; CRS-R = Coma Recovery Scale-Revised; LG = ‘Local-Global’ paradigm.

No effect of ‘Local-Global’ ERP diagnosis was observed on slow-wave density, neither when averaging across the scalp (F_dens_ = 0.68, ^FDR^p_dens_ = 0.51), nor in topographical analyses. By contrast, a significant main effect of functional diagnosis was observed on slow-wave downward slope (F_dslope_ = 3.81, ^FDR^p_dslope_ = 0.02), and patients exhibiting both a local and a global effect showed steeper scalp-averaged downward slopes than patients with no detectable ERP effect (t_dslope(local+global>noeffect)_ = 2.65, ^FDR^p_dslope(local+global>noeffect)_ = 0.026). This effect was further supported by topographical analyses, which revealed a significant main effect of ‘Local-Global’ ERP diagnosis over a broad fronto-central cluster extending towards parietal regions (^CBPS^p_dslope_ < 0.001, 72 electrodes, mass = 289.62). Pairwise contrasts further suggested that this metric tracks the hierarchical level of auditory information integration, most prominently through a frontal cluster differentiating patients exhibiting both a local and a global effect from those showing only a local effect (^CBPS^p_dslope(local+global>local only)_ < 0.001, 19 electrodes, mass = 40.99), thereby linking the global effect, considered a marker of conscious access, to steeper slow-wave downward slopes in frontal regions.

We further examined slow-wave features across ‘Local-Global’ ERP responses within and across CRS-R-defined consciousness states (Supplementary Fig. 2), to determine whether slow-wave metrics could capture neurophysiological functional heterogeneity among patients sharing the same behavioral diagnosis. We then assessed the extent to which this observed functional effect was related to clinical state. Our analyses revealed significant interactions between clinical and functional state at the topographical level, over two clusters for slow-wave density (13 and 15 electrodes; masses = 38.7 and 43.7; both ^CBPS^p_dens_ < 0.001) and three for downward slope (9, 6 and 9 electrodes, masses = 22.8, 16.5 and 28.4, ^CBPS^p_dslope_ < 0.001, 0.004 and < 0.001 respectively). These effects were not apparent when averaging across electrodes (F_dens_ = 1.49, ^FDR^p_dens_ = 0.21, F_dslope_ = 1.73, ^FDR^p_dslope_ = 0.15). These interactions appeared to be primarily driven by differences in slow-wave attributes among patients showing no detectable ERP effect, whose scalp-averaged slow-wave properties varied depending on behavioral state (t_dens(MCS+>UWS)noeffect_ = 2.48, ^FDR^p_dens(MCS+>UWS)noeffect_ = 0.04; t_dslope(MCS+>UWS)noeffect_ = 4.04, ^FDR^p_dslope(MCS+>UWS)noeffect_ < 0.001; t_dslope(MCS->UWS)noeffect_ = 2.71, ^FDR^p_dslope(MCS->UWS)noeffect_ = 0.01), and by a reversal in the direction of the ERP-related contrasts across clinical states: relative to patients with no detectable ERP effect, those showing residual auditory processing displayed higher values in UWS but lower values in MCS+ patients. Most notably, a prominent frontal cluster emerged in MCS+ patients, reflecting steeper downward slopes in patients exhibiting both local and global effects compared with those showing only a local effect (^CBPS^p_dslope(local+global>local only)MCS+_ = 0.002, 6 electrodes, mass = 12.99). No comparable effect was significantly observed in UWS and MCS- patients, suggesting that the relationship between slow-wave intrusion and residual conscious information processing capacities is modulated, at least in part, by the underlying clinical state.

### Slow wave temporal dynamics differ across DoC subgroups

We then explored the temporal dynamics of slow waves throughout the ‘Local-Global’ protocol (∼30 min) to determine whether they reflect progressive changes in brain state over the course of the recording and exhibit signatures of fatigue-related processes (Fig. 3B).

Scalp-averaged slow-wave density evolved linearly across successive ‘Local-Global’ blocks (F_dens_ = 30.90, ^FDR^p_dens_ < 0.001). This effect was significant within each consciousness state, from UWS to MCS+, as well as in healthy controls (t_dens(UWS)_ = 2.80, ^FDR^p_dens(UWS)_ = 0.007; t_dens(MCS-)_ = 2.22, ^FDR^p_dens(MCS-)_ = 0.027; t_dens(MCS+)_ = 3.07, ^FDR^p_dens(MCS+)_ = 0.004; t_dens(CTRL)_ = 3.17, ^FDR^p_dens(CTRL)_ = 0.004), and was associated with widespread scalp clusters in each group (UWS: 108 electrodes, masses = 53.5 and 249.8; MCS-: 82 electrodes, mass = 228.4; MCS+: 127 electrodes, mass = 368.5; CTRL: 131 electrodes, mass = 389.6; all ^CBPS^p_dens_ < 0.001), with an additional small posterior cluster in MCS- patients (3 electrodes, mass = 7.3, ^CBPS^p_dens(MCS-)_ = 0.024). No interaction with behavioral state was observed when disregarding spatial distribution (F_dens_ = 1.31, ^FDR^p_dens_ = 0.27). Consistently, contrasting the first and last blocks of the recording (Supplementary Fig. 3) revealed a significant increase in scalp-averaged slow-wave density in MCS+ patients and healthy controls only (t_dens(End>Start)MCS+_ = 2.15, ^FDR^p_dens(End>Start)MCS+_ = 0.03; t_dens(End>Start)CTRL_ = 2.17, ^FDR^p_dens(End>Start)CTRL_ = 0.03), whereas UWS and MCS- patients showed no detectable change (t_dens(End>Start)MCS-_ = 1.14, ^FDR^p_dens(End>Start)MCS-_ = 0.26; t_dens(End>Start)UWS_ = 1.16, ^FDR^p_dens(End>Start)UWS_ = 0.25). However, post hoc contrasts did not reveal significant differences in the magnitude of this temporal effect between clinical states when considering scalp-averaged density, although a trend toward stronger density increases was observed in MCS+ patients and healthy controls relative to UWS and MCS- patients (t_dens(MCS+>UWS)_ = 1.52, ^FDR^p_dens(MCS+>UWS)_ = 0.35; t_dens(MCS+>MCS-)_ = 1.27, ^FDR^p_dens(MCS+>MCS-)_ = 0.35).

While not significant scalp-wide, channel-wise analyses revealed a significant interaction effect between time and clinical state on slow-wave density, distributed across five circumscribed clusters (3 to 8 electrodes each, 27 in total; cluster masses = 12.0 to 27.4; ^CBPS^p_dens_ ≤ 0.038) over central, temporo-parietal and posterior regions (Fig. 3B). Accordingly, pairwise topographical contrasts revealed significant spatial clusters differentiating consciousness states with respect to the effect of time on slow-wave intrusions. Overall, the magnitude of this temporal effect on slow-wave density appeared to increase with the level of consciousness recovery in specific scalp regions, as evidenced by positive clusters for all successive state comparisons (CBPSp_dens(MCS->UWS)_ < 0.001, 5 electrodes, mass = 11.97; ^CBPS^p_dens(MCS+>UWS)_ ≤ 0.022, three clusters totalling 34 electrodes, masses = 43.5, 7.3 and 28.1; ^CBPS^p_dens(MCS+>MCS-)_ ≤ 0.037, three clusters totalling 20 electrodes, masses = 28.1, 6.8 and 15.5; ^CBPS^p_dens(CTRL>MCS+)_ ≤ 0.003, two clusters totalling 11 electrodes, masses = 17.1 and 10.1), except for two small negative clusters showing the opposite pattern in the MCS->UWS and CTRL>MCS+ contrasts (^CBPS^p_dens(MCS->UWS)_ = 0.049, 3 electrodes, mass = 6.51; ^CBPS^p_dens(CTRL>MCS+)_ = 0.030, 3 electrodes, mass = 7.1).

This state-dependent modulation of temporal dynamics appeared even more pronounced for slow-wave downward slope. Although a significant main effect of time was observed when considering all states together (F_dslope_ = 33.74, ^FDR^p_dslope_ < 0.001), this effect appeared to be primarily driven by MCS+ patients (t_dslope(MCS+)_ = 2.44, ^FDR^p_dslope(MCS+)_ = 0.03) and healthy controls (t_dslope(CTRL)_ = 6.47, ^FDR^p_dslope(CTRL)_ < 0.001). In contrast, no significant scalp-averaged temporal effect was observed on UWS and MCS- patients’ descending slow-wave slopes (t_dslope(MCS-)_ = 0.93, ^FDR^p_dslope(MCS-)_ = 0.47; t_dslope(UWS)_ = 0.62, ^FDR^p_dslope(UWS)_ = 0.53). Both scalp-wide and channel-wise analyses revealed a strong interaction effect between time and consciousness state (F_dslope_ = 10.51, ^FDR^p_dslope_ < 0.001, ^CBPS^p_dslope_ < 0.001, main cluster of 82 electrodes, mass = 410.1; four additional clusters totalling 30 electrodes, ^CBPS^p_dslope_ ≤ 0.028). Consistently, the magnitude and spatial extent of the temporal effect seemed to increase with consciousness level, with widespread positive clusters observed in MCS+ patients and healthy controls (^CBPS^p_dslope(MCS+)_ < 0.001, 62 electrodes, mass = 156.3, plus a small additional cluster of 3 electrodes, mass = 7.0, ^CBPS^p_dslope(MCS+)_ = 0.032 / ^CBPS^p_dslope(CTRL)_ < 0.001, 160 electrodes, mass = 540.5), compared with only small and spatially restricted clusters in UWS and MCS-patients (^CBPS^p_dslope(UWS)_ < 0.001, 8 electrodes, mass = 20.1; ^CBPS^p_dslope(MCS-)_ < 0.001, 8 electrodes, mass = 17.9). Pairwise topographical analyses further revealed marked increases in the temporal modulation of slow-wave downward slope with consciousness recovery (^CBPS^p_dslope(MCS+>UWS)_ ≤ 0.022, five clusters totalling 41 electrodes, masses = 7.4 to 46.3; ^CBPS^p_dslope(MCS+>MCS-)_ < 0.001, two clusters totalling 25 electrodes, masses = 46.9 and 12.2; ^CBPS^p_dslope(CTRL>MCS+)_ ≤ 0.005, five clusters totalling 38 electrodes, masses = 9.6 to 48.7), although this pattern did not extend to the MCS->UWS contrast.

Together, these findings indicate a progressive accumulation of slow waves throughout the ‘Local-Global’ recording, which appears to be, at least spatially, modulated by CRS-R consciousness level. More strikingly, the temporal modulation of slow-wave downward slope seemed to differentiate UWS and MCS- patients from MCS+ patients (and healthy controls).

### Slow waves carry prognostic information

We next examined whether these high-amplitude slow-wave intrusions carried prognostic information by testing their association with long-term consciousness (Fig. 4A) and functional (Fig. 4B) outcomes.

**Figure 4:**
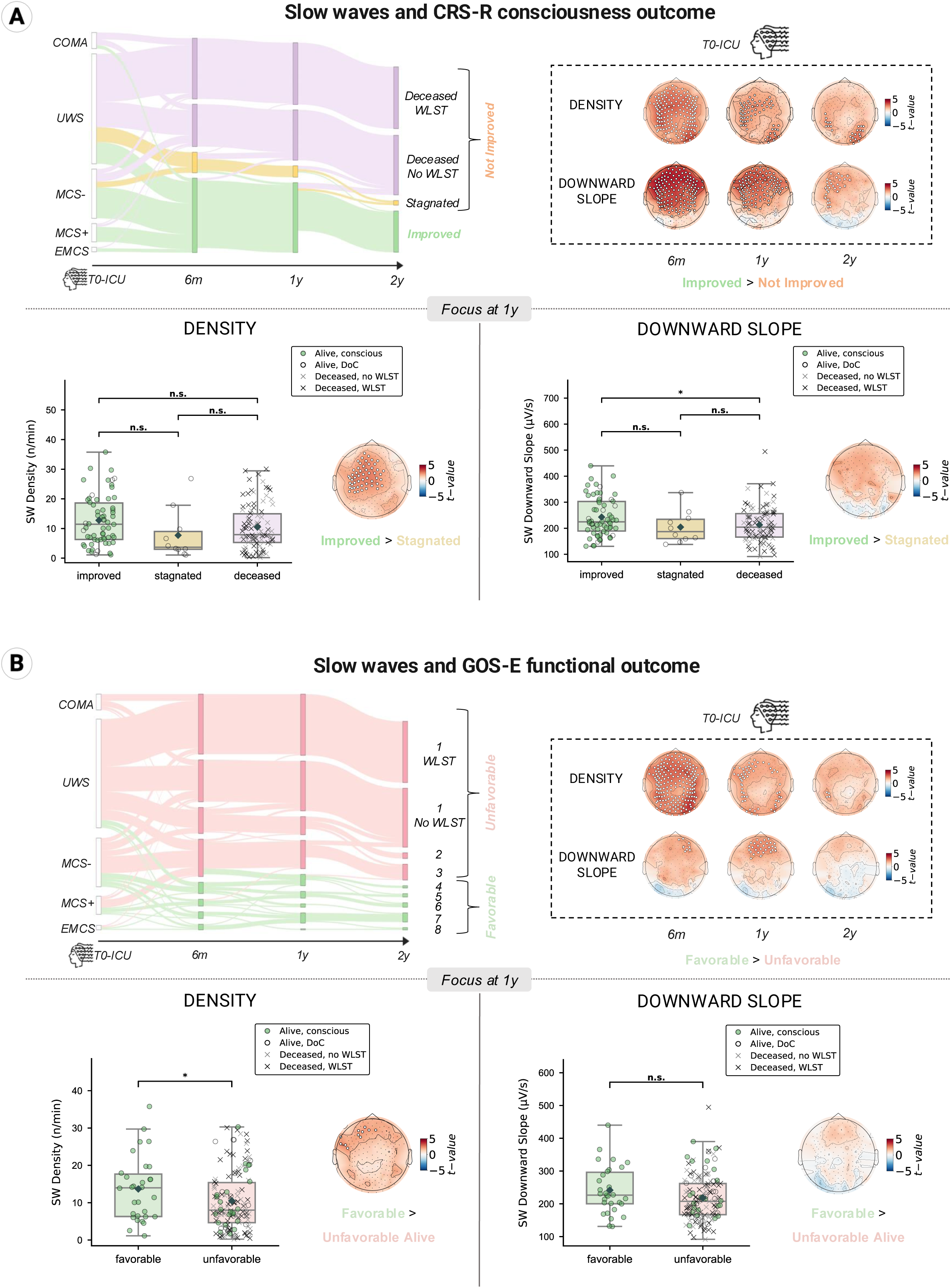
Slow-wave properties predict long-term clinical and functional outcomes. Slow-wave metrics differ according to recovery outcomes assessed at 6 months, 1 year, and 2 years following ICU evaluation using the CRS-R consciousness scale **(A)** and the functional GOS-E scale **(B)**. For CRS-R outcomes (A), “improved” indicates patients whose follow-up consciousness state exceeded the CRS-R state associated with the ICU EEG assessment, whereas “stagnated” indicates patients remaining in the same consciousness state. “Deceased” refers to patients who died either spontaneously or following WLST. The “non-improved” category includes both “stagnated” and “deceased” patients. GOS-E functional outcomes (B), ranging from 1 to 8, were dichotomized into unfavorable (<4) and favorable (≥4) outcomes. Sankey diagrams (top left) illustrate longitudinal transitions in clinical and functional outcomes across follow-up time points, with link widths proportional to the number of patients transitioning between states. Top right panels display statistical topographies of CBPS-corrected t-values for the contrast and slow-wave metric of interest at each follow-up time point (white dot: p < 0.05). Bottom panels further focused on 1 year recovery. Boxplots (left: density; right: downward slope) show the median (central line) and interquartile range (Q1-Q3, box edges), with whiskers extending to the most extreme values within 1.5 × IQR of the box edges. Group-level means are indicated by grey diamonds, while overlaid scatterplots display individual data points. Green-filled circles indicate alive conscious patients, open circles alive DoC patients, grey crosses deceased patients without WLST, and black crosses deceased patients following WLST. Statistical annotations above the boxplots indicate FDR-corrected pairwise differences between outcome groups, derived from linear models (*: p < 0.05, n.s.: non-significant). Scalp topographies displayed to the right of the boxplots illustrate the spatial distribution of CBPS-corrected t-values for the contrast of interest (white dot: p < 0.05). CBPS = cluster-based permutation statistics; CRS-R = Coma Recovery Scale-Revised; GOS-E = Glasgow Outcome Scale-Extended; WLST = Withdrawal of Life Sustaining Therapies.

A Sankey diagram illustrates the evolution of consciousness state between the multimodal ICU assessment when the EEG recording took place and follow-up evaluations at 6 months, 1 year, and 2 years. Relative to their baseline CRS-R-defined state, patients were categorized as ‘improved’, ‘stagnated’, or ‘deceased’, either spontaneously or following withdrawal of life-sustaining therapies (WLST). This visualization revealed substantial heterogeneity in recovery trajectories. Unsurprisingly, patients with higher baseline levels of consciousness had more favorable outcomes.

Slow waves could explain some of the within- and across-group differences. Topographical contrast analyses comparing baseline slow-wave profile of patients who subsequently showed consciousness improvement with those who did not (including both stagnated and deceased outcomes) revealed significant clusters of increased slow-wave density and steeper downward slopes, for all three follow-up stages (Fig. 4A; 6 months: ^CBPS^p_dens(Impr>NotImpr)6m_ < 0.001, 162 electrodes, mass = 417.12; ^CBPS^p_dslope(Impr>NotImpr)6m_ < 0.001, 136 electrodes, mass = 463.14; 1 year: ^CBPS^p_dens(Impr>NotImpr)1y_ < 0.001, two clusters totalling 65 electrodes, masses = 111.38 and 32.65; ^CBPS^p_dslope(Impr>NotImpr)1y_ < 0.001, 108 electrodes, mass = 289.49; 2 years: ^CBPS^p_dens(Impr>NotImpr)2y_ < 0.001, two clusters totalling 25 electrodes, masses = 19.84 and 38.23; ^CBPS^p_dslope(Impr>NotImpr)2y_ < 0.001, two clusters totalling 30 electrodes, masses = 54.39 and 11.06). These effects remained essentially unchanged after excluding patients who died following WLST (Supplementary Fig. 4).

To better isolate markers of consciousness recovery, we then excluded deceased patients and focused on the comparison between ‘improved’ and ‘stagnated’ outcomes at the 1-year follow-up, a clinically relevant time point capturing both the relatively rapid recovery trajectory of post-anoxic patients and the often-slower recovery observed following traumatic brain injury^91^. Topographical analyses revealed a large fronto-central positive cluster (^CBPS^p_dens(Impr>Stag)1y_ < 0.001, 53 electrodes, mass = 119.15), indicating that patients whose consciousness state improved over the following year exhibited markedly higher slow-wave density within this region at T0 than patients whose consciousness state did not evolve. However, no significant difference in scalp-averaged slow-wave density was observed (t_dens(Impr>Stag)1y_ = 1.86, ^FDR^p_dens(Impr>Stag)1y_ = 0.12). Likewise, slow-wave downward slope did not differ between ‘improved’ and ‘stagnated’ outcome groups at 1 year, neither in scalp-averaged analyses (t_dslope(Impr>Stag)1y_ = 1.63, ^FDR^p_dslope(Impr>Stag)1y_ = 0.16) nor in channel-wise topographical analyses, suggesting that its prognostic value was mainly driven by mortality.

We further refined these analyses by focusing on functional recovery (GOS-E Sankey diagrams; favorable outcome: ≥4; unfavorable outcome: <4; see Methods for details). Channel-wise analyses comparing baseline slow-wave density between patients who subsequently achieved favorable and unfavorable outcomes at 6 months, 1 year, and 2 years after ICU assessment revealed positive clusters, indicating increased slow-wave intrusions in patients with favorable long-term recovery. This effect was particularly pronounced and spatially widespread when predicting 6-month outcomes (^CBPS^pdens(Fav>Unfav)6m < 0.001, 178 electrodes, mass = 484.39), but remained detectable at 1 year over frontal, temporal, and posterior regions (^CBPS^pdens(Fav>Unfav)1y < 0.001, three clusters totalling 59 electrodes, masses = 86.93, 15.65 and 24.95). At 2 years, the effect became more spatially restricted, where only a small left temporal cluster survived correction for multiple comparisons (^CBPS^pdens(Fav>Unfav)2y = 0.012, 4 electrodes, mass = 8.33). Interestingly, downward slope predicted 1-year functional outcome over a frontal cluster (^CBPS^p_dslope_(Fav>Unfav)1y < 0.001, 29 electrodes, mass = 65.42), overlapping with the region where steeper slopes were associated with the presence of a global effect (Fig. 3A).

Consistent with these topographical findings, patients with favorable outcomes at 1 year showed increased scalp-averaged baseline slow-wave density compared to patients with unfavorable outcomes (t_dens(Fav>Unfav)1y_ = 1.98, ^FDR^p_dens(Fav>Unfav)1y_ = 0.049), whereas no significant group difference was observed for downward slope (t_dslope(Fav>Unfav)1y_ = 1.67, ^FDR^p_dslope(Fav>Unfav)1y_ = 0.097). To further disentangle the contribution of mortality to these effects, we restricted the analysis to surviving patients and compared favorable outcomes with ‘unfavorable-alive’ outcomes, comprising conscious patients with severe disability and patients who remained in a DoC. Despite a markedly reduced sample size, a significant anterior positive cluster was observed for slow-wave density (^CBPS^p_dens(Fav>UnfavAlive)1y_ < 0.001, 7 electrodes, mass = 14.8; second cluster: 0.006, 4 electrodes, mass = 9.2), indicating that enhanced slow-wave intrusions within frontal regions were specifically associated with subsequent recovery of functional independence rather than survival alone. By contrast, no cluster survived correction for descending slope.

Together, these findings suggest that slow-wave activity measured during the acute phase may carry clinically relevant prognostic information and be associated with both long-term recovery of consciousness and functional independence.

### Slow waves correlate with structural brain integrity

Given the associations observed between slow-wave properties and both diagnostic and prognostic indices of consciousness, we next sought to determine whether these neurophysiological signatures were linked to the preservation of structural brain integrity.

We first investigated the relationship between slow-wave properties and fractional anisotropy derived from diffusion MRI (Fig. 5A), an indirect marker of white-matter structural integrity (see Methods). Our analyses focused on projection fibers, which support communication between cortical and subcortical regions. We observed a positive association between projection fibers FA and scalp-averaged slow-wave downward slope across the combined cohort of UWS, MCS-, and MCS+ patients (_ρdslope_ = 0.27, _pdslope_ = 0.011). Projection fibers FA itself did not differ between the two major etiological groups (EMM_Anox_ = 0.734, EMM_TBI_ = 0.782, t_FA(TBI>Anox)_ = 1.30; ^FDR^p_FA(TBI>Anox)_ = 0.20) (Supplementary Fig. 5). We nonetheless examined this FA-slow-wave relationship separately within post-traumatic and post-anoxic patients, given their distinct underlying pathophysiological mechanisms. Higher projection fibers FA was associated with steeper slow-wave downward slopes in both etiological groups (TBI: _ρdslope(TBI)_ = 0.60, ^FDR^p_dslope(TBI)_ = 0.027; Anoxia: _ρdslope(Anox)_ = 0.30, ^FDR^p_dslope(Anox)_ = 0.045). Although the coefficient was numerically larger in the TBI group, the two correlations did not differ significantly (Fisher’s z = 1.17, p = 0.24). Similar results were obtained after adjustment for age using partial Spearman correlations (partial _ρdslope(TBI)_ = 0.60, ^FDR^p_dslope (TBI)_ = 0.023; partial _ρdslope(Anox)_ = 0.35, ^FDR^p_dslope(Anox)_ = 0.023), indicating that these associations were not accounted for by age differences between patients.

**Figure 5:**
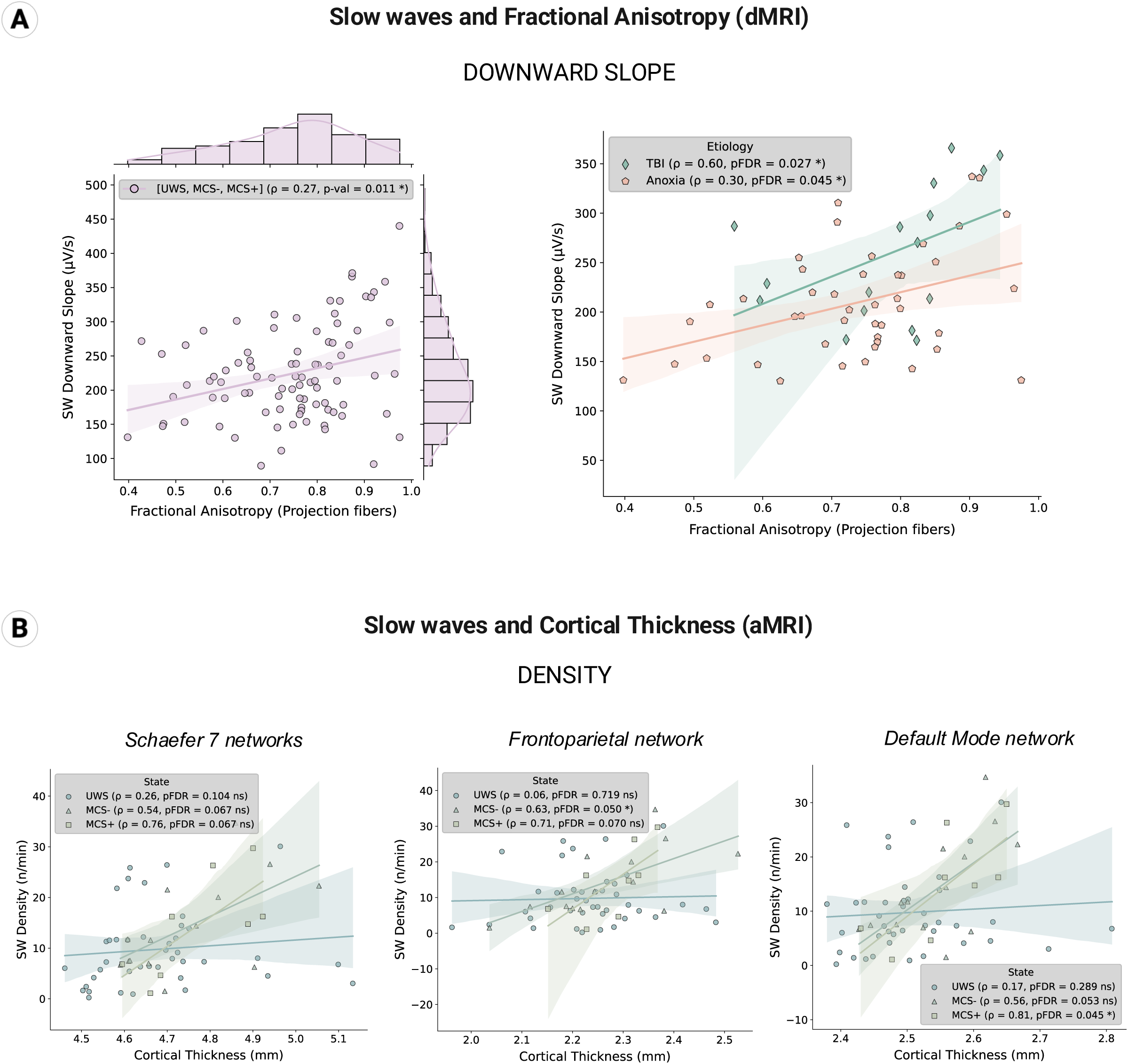
Slow-wave properties are associated with markers of structural brain integrity. **(A)** Slow-wave downward slope is positively correlated to Projection fibers fractional anisotropy across the whole cohort of UWS, MCS- and MCS+ patients (left) and stratified by etiology (TBI/Anoxia; right). **(B)** Slow-wave density is associated with cortical thickness within specific clinical states and functional networks. (A-B) Correlations were assessed using Spearman rank correlations. Scatterplots display subject-level associations between slow wave properties and the corresponding MRI metrics, with regression lines shown for visualization. Spearman correlation coefficients (ρ) and associated FDR-corrected p-values are displayed for each analysis, with asterisks indicating significance levels (*: p < 0.05, n.s.: non-significant). aMRI = anatomical Magnetic Resonance Imaging; dMRI = diffusion Magnetic Resonance Imaging/DTI = Diffusion Tensor Imaging; FA = Fractional Anisotropy; ROI = Region of Interest; TBI = Traumatic Brain Injury.

We further assessed the association between high-amplitude slow-wave intrusions and cortical thickness (Fig. 5B), an index of cortical structural integrity, within several large-scale functional networks. Cortical thickness did not differ significantly between CRS-R states in the frontoparietal (F_CortThick_ =1.26, ^FDR^p_CortThick_ = 0.30) or default mode networks (F_CortThick_ = 1.42, ^FDR^p_CortThick_ = 0.26) (Supplementary Fig. 5). Across the Schaefer-Yeo 7-networks parcellation, thickness tended to increase with the level of consciousness recovery, although this effect did not reach the corrected threshold (F_CortThick_ = 3.17, ^FDR^p_CortThick_ = 0.054) and no pairwise contrast was significant after FDR correction. Group-level differences in cortical structure were therefore modest, and all correlations reported below were computed within CRS-R groups. Positive associations between cortical thickness and slow-wave density were observed across networks and in both MCS subgroups. Across Schaefer-Yeo 7-networks, coefficients were moderate to strong in MCS- and MCS+ patients but did not survive FDR correction (ρ_dens (MCS-)All7_ = 0.54, ^FDR^p_dens (MCS-)All7_ = 0.067; ρ_dens (MCS+)All7_ = 0.76, ^FDR^p_dens (MCS+)All7_ = 0.067). However, within the frontoparietal network, the association was significant in MCS- patients (ρ_dens(MCS-)FPN_ = 0.63, ^FDR^p_dens(MCS-)FPN_ = 0.050), with a numerically larger coefficient in MCS+ patients falling just short of it (ρ_dens(MCS+)FPN_ = 0.71, ^FDR^p_dens(MCS+)FPN_ = 0.070); this discrepancy may reflect the smaller size of the MCS+ subgroup rather than a weaker association. A comparable pattern was observed within the default mode network, where the correlation reached significance in MCS+ patients (ρ_dens(MCS+)DMN_ = 0.81, ^FDR^p_dens(MCS+)DMN_ = 0.045), and remained marginal in MCS- patients (ρ_dens(MCS-)DMN_ = 0.56, ^FDR^p_dens (MCS-)DMN_ = 0.053). No such association was detected in UWS patients in any of those three networks (all ^FDR^p > 0.10).

Overall, these results support a relationship between high-amplitude slow-wave expression and the preservation of structural brain integrity, encompassing both cortico-subcortical white-matter pathways and cortical networks, the latter showing the strongest associations in MCS states.

### Single-patient classification from slow-wave features

To extend these group-level findings, we next investigated whether combining multiple slow-wave features could provide clinically relevant patient-level diagnostic and prognostic predictions.

Using a multivariate Partial Least Squares Discriminant Analysis framework (PLS-DA, see Methods), we evaluated the ability of slow-wave metrics to discriminate consciousness states at the time of assessment (Fig. 6A; CRS-R diagnosis) and to predict subsequent improvement of consciousness (Fig. 6B left; Improved versus Not Improved) and functional outcome (Fig. 6B right; Favorable versus Unfavorable; Fig. 4B).

**Figure 6:**
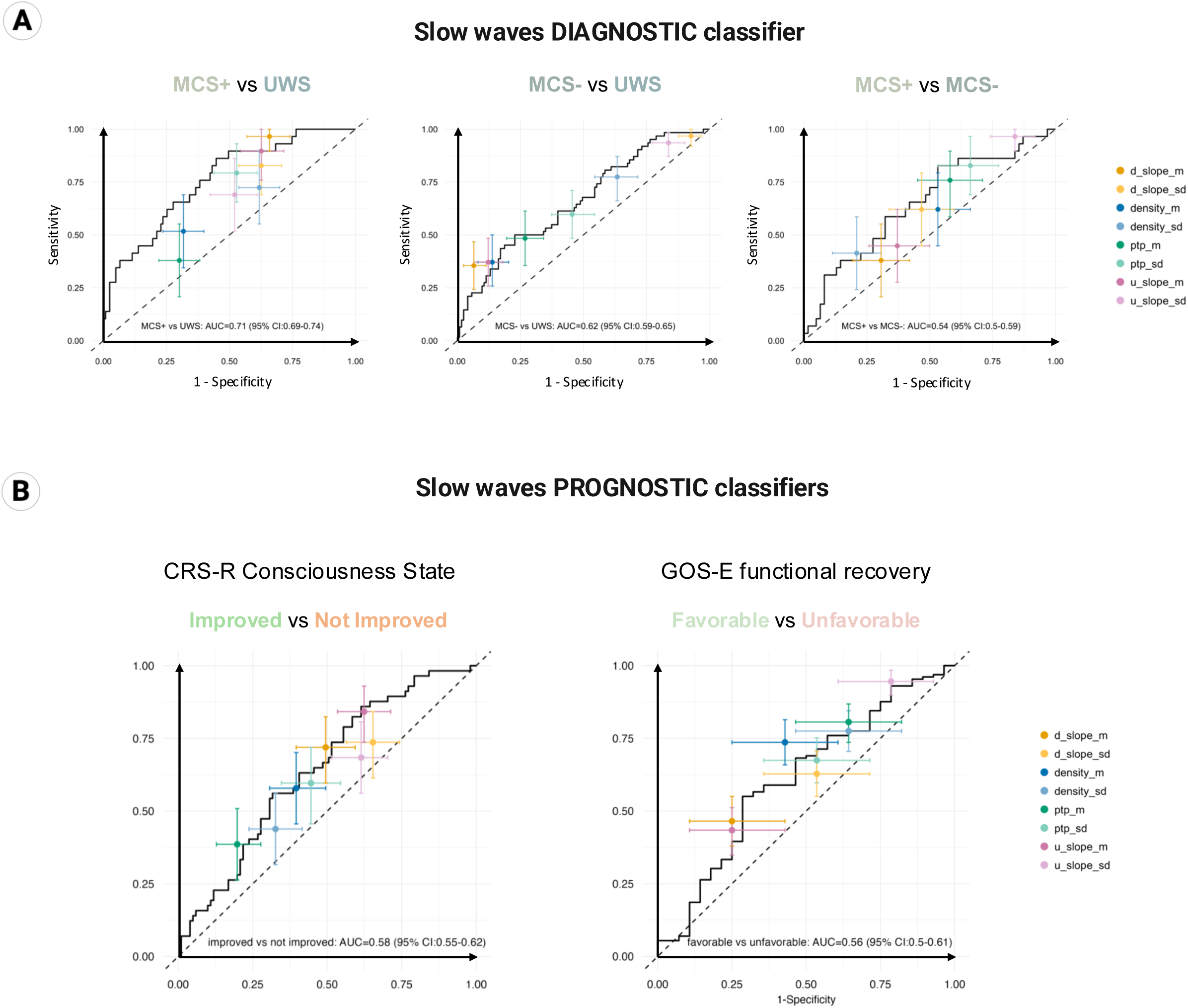
Single-patient classification of diagnosis and outcome from multivariate slow-wave features. Receiver operating characteristic (ROC) curves for multivariate PLS-DA models discriminating CRS-R groups **(A)** and one-year outcomes **(B)** (analyses restricted to patients recorded in the acute phase). Black curves show the ROC of each multivariate classifier, computed on the full sample and shown for illustration; the cross-validated area under the curve (AUC) reported within each panel was obtained by 5-fold cross-validation with 20 repetitions and is reported as the mean across repetitions ±1.96 SD. Colored dots indicate the operating point of each individual slow-wave feature at its optimal threshold (Youden index), with horizontal and vertical bars representing bootstrap 95% confidence intervals for specificity and sensitivity, respectively (2,000 resamples). These univariate operating points were determined on the same sample (training dataset) and are therefore optimistic; they are shown to compare the profiles of individual features rather than to estimate standalone diagnostic performance. For each feature, ‘m’ denotes the average across all EEG channels and recording blocks, whereas ‘sd’ denotes the standard deviation across the eight recording blocks after averaging across channels, reflecting temporal variability. AUC = Area Under the Curve; CI = Confidence Interval; d_slope = Downward Slope; ptp = Peak-to-Peak Amplitude; u_slope = Upward Slope.

For diagnostic classification (Fig. 6A), the best performance was obtained for the MCS+ versus UWS contrast (cross-validated AUC = 0.71 [0.69-0.74], mean ± 1.96 × SD across repeated cross-validation runs). Performance was lower for the MCS- versus UWS contrast (cross-validated AUC = 0.62 [0.59-0.65]) and did not exceed chance level for MCS- versus MCS+ (cross-validated AUC = 0.54 [0.50-0.59]). This ordering, with discrimination improving as the clinical distance between states increases, supports the construct validity of slow-wave metrics as markers of consciousness recovery, while indicating that they do not capture the MCS-/MCS+ distinction.

While combining slow-wave features appeared informative for discriminating the current level of consciousness, it showed little predictive value for one-year outcome. Cross-validated AUCs reached 0.58 [0.55-0.62] for subsequent improvement of consciousness (Improved versus Not Improved) and 0.56 [0.50-0.61] for functional outcome at one year (Favorable versus Unfavorable), above chance for the former but below the range typically considered clinically actionable. Among individual metrics, mean slow-wave density showed the most balanced profile (sensitivity 0.74 [0.66-0.81], specificity 0.57 [0.39-0.75]), while features reaching higher sensitivity, such as upward slope temporal variability, did so at the cost of near-absent specificity (sensitivity 0.95 [0.90-0.98], specificity 0.21 [0.07-0.39]), consistent with the limited overall performance of the prognostic models.

Importantly, classifiers’ performance was essentially unchanged when patients who died following WLST were excluded (Supplementary Fig. 6; AUC = 0.58 [0.53-0.63] and 0.57 [0.53-0.62] for improvement of consciousness and functional outcome, respectively), indicating that classifier performance was not driven by self-fulfilling prophecy.

## DISCUSSION

### A single EEG signature indexing multiple dimensions of consciousness recovery

Collectively, our findings establish spontaneous slow-wave dynamics as a versatile electrophysiological marker of consciousness recovery following severe brain injury. Estimated from a single non-invasive EEG recording, it correlates with indices of behavioral responsiveness, residual conscious auditory processing, and long-term consciousness outcome.

With respect to diagnosis, slow-wave dynamics tracked richer states of consciousness as defined through behavioral or neurophysiological indices. Slow-wave density and downward slope were both higher in MCS- than UWS patients over a diffuse scalp cluster, consistent with their greater behavioral responsiveness as assessed with the CRS-R (Fig. 2A). This effect held true at the single-patient level: a classifier (Fig. 6A) built with slow-wave features achieved above-chance performance in predicting if the patient was in a UWS or MCS- state, showing high promise for helping to disambiguate these two clinically close states. Additionally, region-specific increases in slow-wave downward slope mirrored improvements in auditory information processing capacity (Fig. 3A), as assessed with the ‘Local-Global’ ERP paradigm^25^. Steeper anterior descending slopes were associated with the presence of a ‘global effect’, a P3b thought to be a neural marker of conscious access^92^, echoing the prominent frontal role attributed to conscious access by the Global Neuronal Workspace and Higher-Order Theory frameworks^93–97^.

With respect to prognosis, higher slow-wave density and steeper downward slopes predicted improvements of consciousness (Fig. 4A) and more favorable functional outcomes on the GOS-E (Fig. 4B) over time, with the most widespread effects observed at six months and strongly persisting at one year. Slow-wave density further distinguished patients who ultimately recovered functional independence from those surviving with persistent DoC or severe disability. These associations remained essentially unchanged after excluding patients who died following withdrawal of life-sustaining therapy (Supplementary Fig. 4), arguing against a self-fulfilling prophecy bias^98^. However, while diagnostic classification proved promising at the single-patient level, single-patient prognostic classification remained modest (Fig. 6B), suggesting that further refinement, potentially through the integration of spatial variability features, will be needed to translate these group-level associations into individual clinical predictions.

### A common electrophysiological phenotype arising from multiple mechanisms

This diagnostic and prognostic versatility suggests that spontaneous slow-wave properties capture a diverse set of structural and functional aspects of brain physiology^14^, shaped both by the integrity of large-scale cortico-subcortical networks and by the dynamic modulation of their physiological state.

Slow-wave dynamics appeared closely tied to the preservation of cortico-subcortical networks (Fig. 5). Our results revealed a markedly non-linear relationship between slow-wave dynamics and CRS-R-defined levels of consciousness (Fig. 2A): most slow-wave features followed an inverted U-shaped profile, peaking in MCS patients while remaining lower in UWS, suggesting that progressive disruption of network integrity paradoxically reduces slow-wave expression despite greater clinical impairment, a pattern at odds with the classical association between cortical slowing and neurological insult^43–45,99,100^. Consistent with this, slow-wave density was positively associated with cortical thickness in consciousness-related networks (frontoparietal and default mode^101^) in MCS states (Fig. 5B), and slow-wave downward slope was positively associated with fractional anisotropy within projection fibers (Fig. 5A), suggesting that preserved subcortico-cortical connectivity, including thalamo-cortical projections, supports the large-scale synchronization underlying slow-wave generation. Together, these findings converge with current models (the mesocircuit hypothesis and the ABCD model^2,69,102,103^) and theories of consciousness such as the Global Neuronal Workspace^94^ emphasizing the central role of long-range cortico-cortical and thalamo-cortical circuitry in consciousness recovery.

If spontaneous slow-wave generation depends on preserved structural architecture, differences in brain damage pattern across etiologies should shape their expression. Indeed, post-traumatic patients exhibited higher slow-wave density and steeper downward slopes than post-anoxic patients, particularly at lower levels of consciousness (Fig. 2B). This likely reflects the diffuse cortico-subcortical disruption typical of anoxic injury, consistent with EEG observations that these patients often show low-voltage activity rather than the cortical slowing typical of TBI^104,105^, and may partly explain the inverted U-shape seen in the overall cohort (Fig. 2A; Supplementary Fig. 1). The strong correlation between downward slope and fractional anisotropy in TBI patients (Fig. 5A) supports a close link to white-matter integrity in this population, though neuroinflammatory processes, which play a prominent and protracted role after TBI^106,107^, may also contribute to the etiological difference.

Several converging pathological processes are thought to promote slow-wave generation after brain injury, largely by disrupting the tonic cortical drive that normally prevents cortical bistability: the alternation between depolarized and hyperpolarized states well characterized in sleep^108,109^ (see Massimini et al., 2024^50^ for a review). These include local neuroinflammation, lesion-associated edema, and deafferentation-driven network disconnection, all of which shift cortical networks toward the inhibitory dominance that favors bistability^110–112^.

Beyond these structural and lesion-related determinants, our findings indicate that slow-wave activity also captures a genuine dynamic functional component reflecting the homeostatic regulation of network physiological states. In the healthy brain, large-amplitude slow waves emerge during physiological sleep and accumulate locally during prolonged wakefulness or sustained cognitive engagement through use-dependent homeostatic regulation^32,34–38^. A similar phenomenon appeared here: slow-wave activity showed progressive within-session temporal modulation whose magnitude and spatial extent varied with clinical level of consciousness (Fig. 3B), being more pronounced in MCS+ patients and healthy controls who alone showed a clear scalp-averaged increase in slow-wave intrusions across the ∼30-minute recording (Supplementary Fig. 3) while less responsive patients showed more limited time-dependent accumulation. This greater ‘fatigue-like’ modulation in patients with behavioral evidence of consciousness may reflect a preserved capacity of cortical networks to adapt dynamically over time, consistent with transient shifts in network configuration favoring conscious processing^113^ and with the GNW view that sustained long-range network mobilization is required for effortful cognitive engagement^94,95^. The prolonged activation of the GNW during the “Local-Global” task could indeed lead to fatigue, possibly expressed in the form of wake slow waves^114^.

While temporal variability in slow-wave features distinguished consciousness states at the group level (Fig. 3B), its contribution at the single-patient level (Fig. 6) was more nuanced: it captured complementary aspects of slow-wave dynamics, favoring sensitivity over specificity, although averaged metrics carried most of the discriminative weight. Its contribution to personalized medicine approaches therefore deserves further investigation, particularly in light of prior evidence that temporal fluctuations in neural or physiological signals carry diagnostic and prognostic value in DoC more broadly, from EEG variability^26,31,115–117^ and olfactory habituation^118^ to fMRI network reconfiguration^28^. Should this hold, temporal variability in slow-wave activity may index the homeostatic regulation of network dynamics, offering a finer-grained marker of network functional efficacy than time-averaged measures alone.

Rather than reflecting a single mechanism, slow-wave dynamics in DoC patients may thus constitute a common electrophysiological phenotype onto which distinct but overlapping pathophysiological processes converge: a uniquely sensitive marker yet intrinsically challenging to interpret mechanistically.

### Slow Waves, Friend or Foe?

Although delta rhythms have traditionally been associated with diminished consciousness^119–121^, and prominent slow waves thought to reflect neuronal OFF periods^89,108,109^, slow-wave expression in the present study consistently predicted richer behavioral responsiveness, stronger residual cognitive processing, and more favorable long-term recovery. We propose this apparent contradiction arises because high-amplitude slow waves are not simply markers of a propensity for cortical bistability, whether sleep-related or injury-induced: their generation also requires sufficient structural and functional preservation of cortico-subcortical networks to synchronize neuronal up- and down-states across distributed populations.

UWS patients show the largest TMS-evoked slow waves^70^, suggesting strong bistability propensity, but the severe network disruption in this population, particularly after anoxic injury, may limit their ability to generate spontaneous slow waves. MCS patients, by contrast, may retain both greater bistability propensity (reflected in their higher slow-wave expression relative to EMCS patients and controls) and sufficient network preservation for these events to emerge spontaneously. We therefore argue that prominent spontaneous slow waves are best viewed not as intrinsically beneficial or detrimental, but as an emergent signature of a brain still capable of generating large-scale coordinated events despite severe injury. This view is supported by previous work on EEG reactivity in DoC patients, stressing that any kind of reactivity should be considered a positive outcome^68^. Relatedly, during healthy sleep, the occurrence of a large SW (e.g., a K-complex) following sensory stimulation is thought to reflect a complex balance between sensory processing and the suppression of this processing^122^.

This interpretation does not preclude a functional contribution of slow waves themselves. As in physiological sleep, where slow waves are thought to support synaptic homeostasis^123–125^, memory consolidation^126–128^, and metabolic clearance^129^, spontaneous slow waves in injured brains might similarly contribute to brain homeostasis^42,50,130^. However, whether slow-wave dynamics in DoC are merely markers of preserved neural integrity or active contributors to the recovery of consciousness remains a key open question.

### Multidimensional slow-wave phenotyping for diagnosis and prognosis

The present study demonstrates that characterizing slow waves at the level of individual events, rather than relying solely on time-averaged spectral slowing, provides clinically meaningful diagnostic and prognostic information by capturing the spatial, temporal, and morphological properties of slow waves. Combining these descriptors yielded promising subject-level classifiers, highlighting the value of multidimensional slow-wave cartography for individualized assessment.

Distinct features may capture complementary physiology: slow-wave density could primarily reflect the propensity of cortical populations to shift to a default state, while downward slope may more specifically index the efficiency of large-scale synchronization during transitions to OFF periods^80,131–134^. These interpretations remain to be directly tested, as correlational analyses cannot disentangle the causal contributions of bistability, network integrity, and homeostatic regulation. Addressing this will require multimodal and interventional approaches combining spontaneous EEG with structural imaging, targeted neuromodulation, and computational modeling.

### Future directions: sleep physiology and therapeutic potential

Our findings highlight the need to better characterize sleep physiology in DoC. If slow-wave expression is partly shaped by sleep homeostasis, as proposed here, its interpretation cannot be dissociated from the profound alterations of sleep architecture, vigilance, and brainstem arousal systems that characterize these patients^135,136^. Sleep spindles have similarly shown diagnostic and prognostic value in predicting consciousness recovery^137^, likely reflecting preserved thalamo-cortical circuitry; investigating how slow waves interact with these signatures, together with PSG measures, may help disentangle their pathophysiological determinants.

If slow-wave dynamics are not only correlates of preserved network integrity but also active contributors to physiological regulation, approaches aimed at modulating them, including pharmacological, sensory, and non-invasive brain stimulation methods already under investigation in sleep research^138–140^, may eventually inspire therapeutic strategies for DoC^50^. Whether such interventions could promote consciousness recovery remains speculative but represents an intriguing avenue for future work.

## SUPPLEMENTARY FIGURES

**Figure S1:**
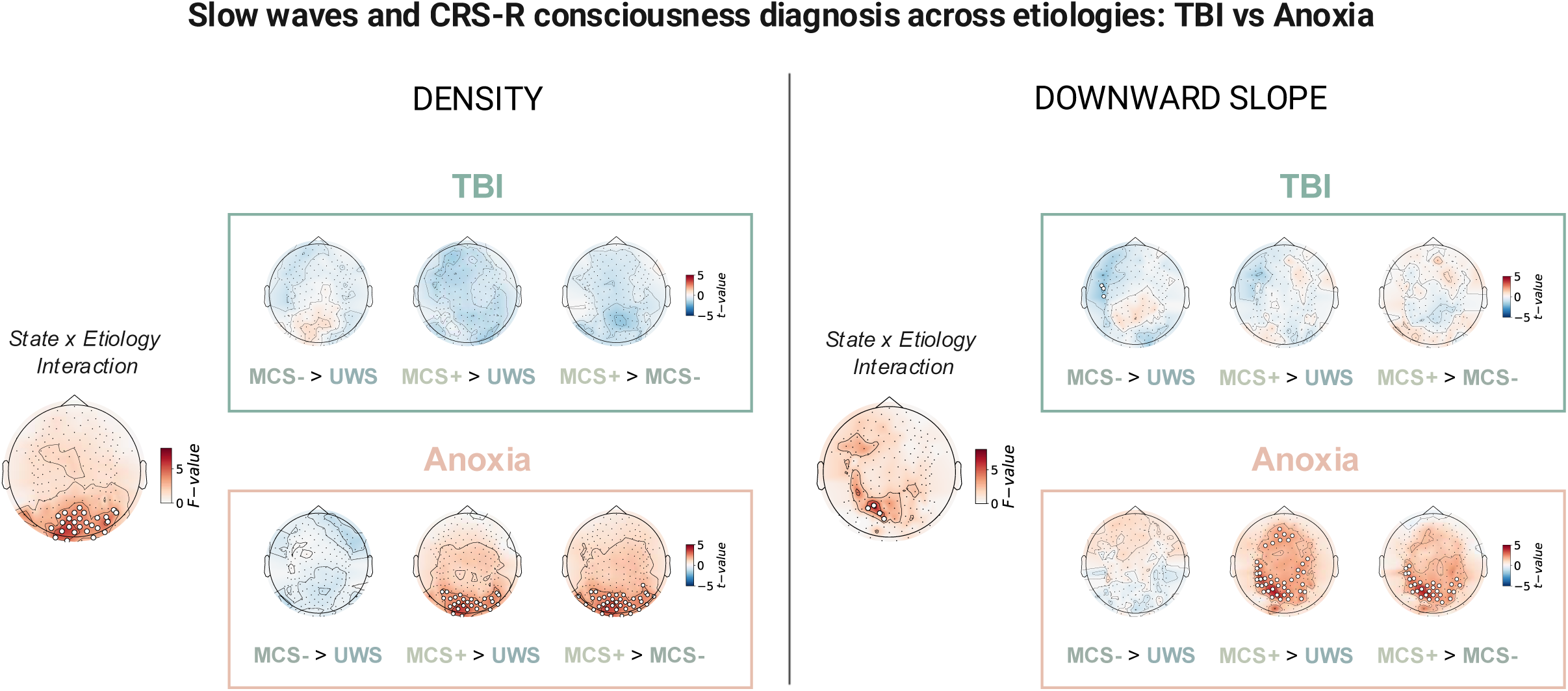
Slow waves and CRS-R consciousness diagnosis across etiologies: TBI vs Anoxia. Scalp topographies of the effect of CRS-R group on slow wave Density (left) and Downward Slope (right) split by etiology: (*top*) TBI patients and (*bottom*) Anoxia patients. Left topographies display statistical topographies of F-values (State * Etiology interaction) and boxed topographies show the t-values by contrast (MCS- > UWS; MCS+ > UWS; MCS+ > MCS-). Channels showing statistically significant effects after CBPS correction (p < 0.05) are marked with white dots.

**Figure S2:**
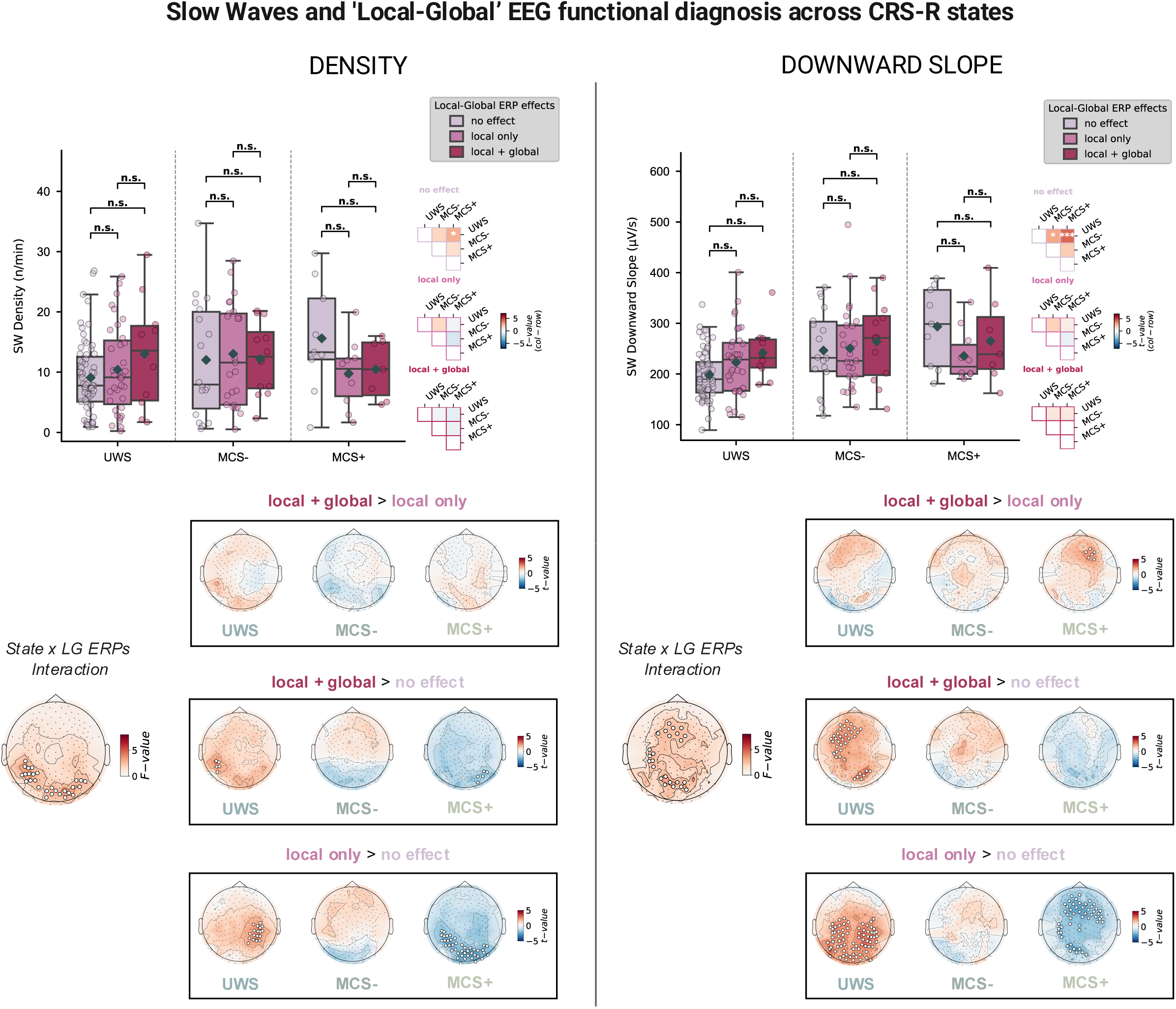
Slow Waves and ‘Local-Global’ EEG functional diagnosis across CRS-R states. Distribution of slow wave Density (left) and Downward Slope (right), averaged across all channels for the patients showing no effect in the ‘Local-Global’ paradigm, only a local effect or both the local and global effect, for USW, MCS- and MCS+ groups separately. Boxplots show the median (central line) and interquartile range (Q1-Q3, box edges), with whiskers extending to the most extreme values within 1.5 × IQR of the box edges. Group-level means are indicated by grey diamonds, while overlaid scatterplots display individual data points (circles). Boxplots are arranged according to increasing levels of consciousness recovery in DoC patients: UWS, MCS-, and MCS+. The triangle matrices display pairwise comparisons between states for the different functional groups, derived from linear models. Colors represent the t-statistics associated with each contrast (column - row), while asterisks indicate FDR-corrected significance levels (*: p < 0.05, **: p < 0.01, ***: p < 0.001). Asterisks above the boxplots indicate FDR-corrected between-etiology differences within each state, using the same significance thresholds; n.s., non-significant. Scalp topographies displayed below illustrate the spatial distribution of the F-values testing the State * LG effect (no effect; local only; local + global; left topographies) and t-values contrasting the different functional groups by CRS-R state (boxed topographies). Channels showing statistically significant effects after CBPS correction (p < 0.05) are marked with white dots.

**Figure S3:**
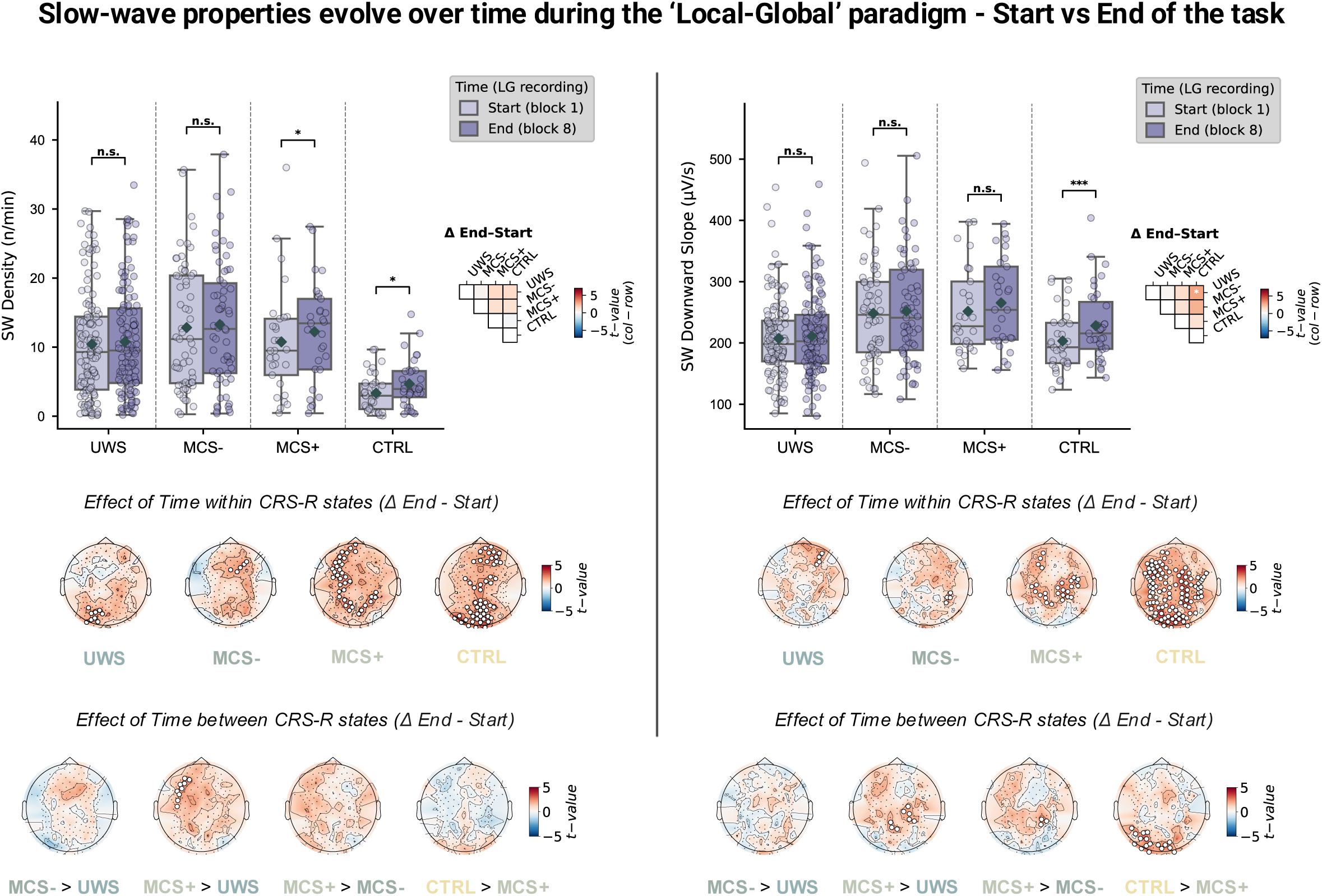
Slow-wave properties evolve over time during the ‘Local-Global’ paradigm - Start vs End of the task. Distribution of slow wave Density (left) and Downward Slope (right), averaged across all channels at the beginning (1^st^ block) and end (last block) of the ‘Local-Global’ paradigm for each consciousness state. Boxplots show the median (central line) and interquartile range (Q1-Q3, box edges), with whiskers extending to the most extreme values within 1.5 × IQR of the box edges. Group-level means are indicated by grey diamonds, while overlaid scatterplots display individual data points (circles). Boxplots are arranged according to increasing levels of consciousness recovery in DoC patients: UWS, MCS-, MCS+ and CTRL. CTRL refers to healthy controls. The triangle matrix displays pairwise comparisons between states, derived from linear models. Colors represent the t-statistics associated with each contrast (column - row), while asterisks indicate FDR-corrected significance levels (*: p < 0.05, **: p < 0.01, ***: p < 0.001). Asterisks above the boxplots indicate FDR-corrected between-etiology differences within each state, using the same significance thresholds; n.s., non-significant. Scalp topographies displayed below illustrate the spatial distribution of the t-values contrasting the last and first blocks within (top) and between CRS-R states (bottom). Channels showing statistically significant effects after CBPS correction (p < 0.05) are marked with white dots.

**Figure S4:**
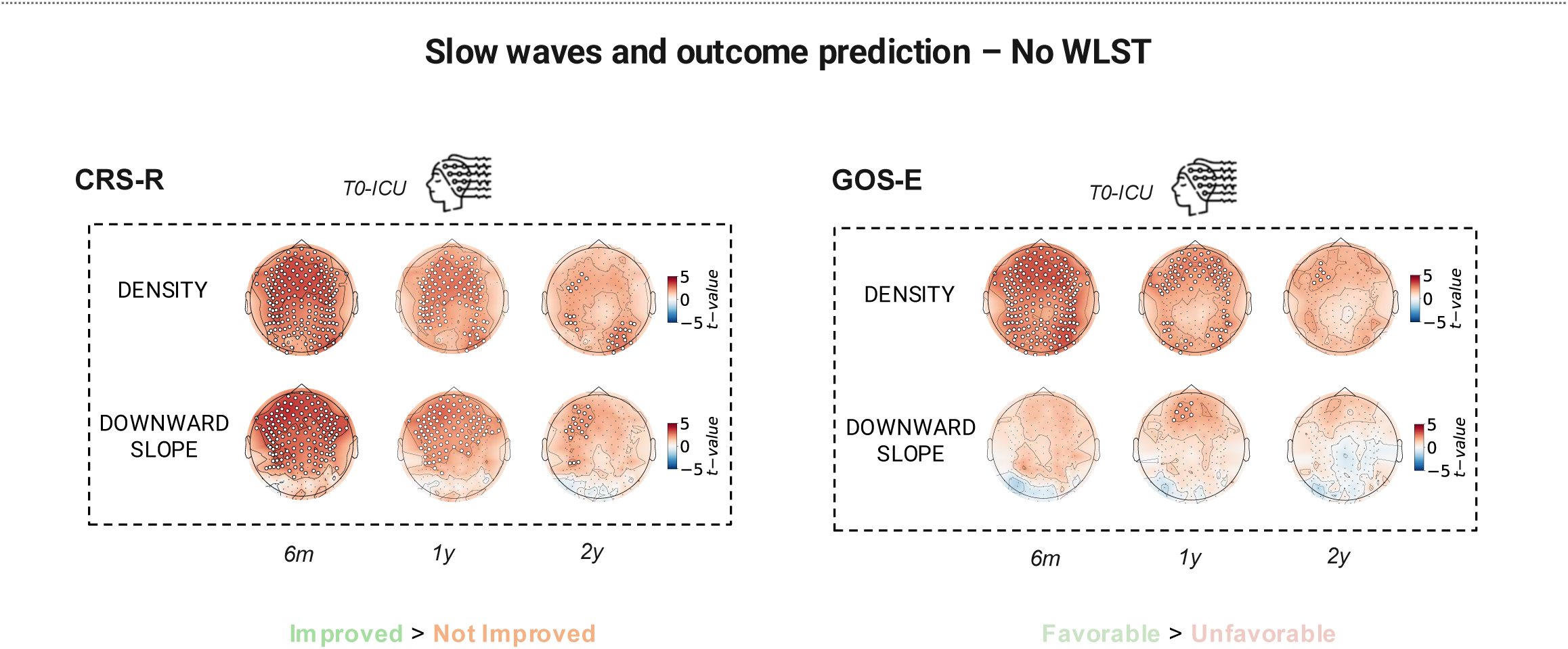
Slow waves and outcome prediction – No WLST. Same as Figure 4A right and 4B left but after removing WLST patients.

**Figure S5:**
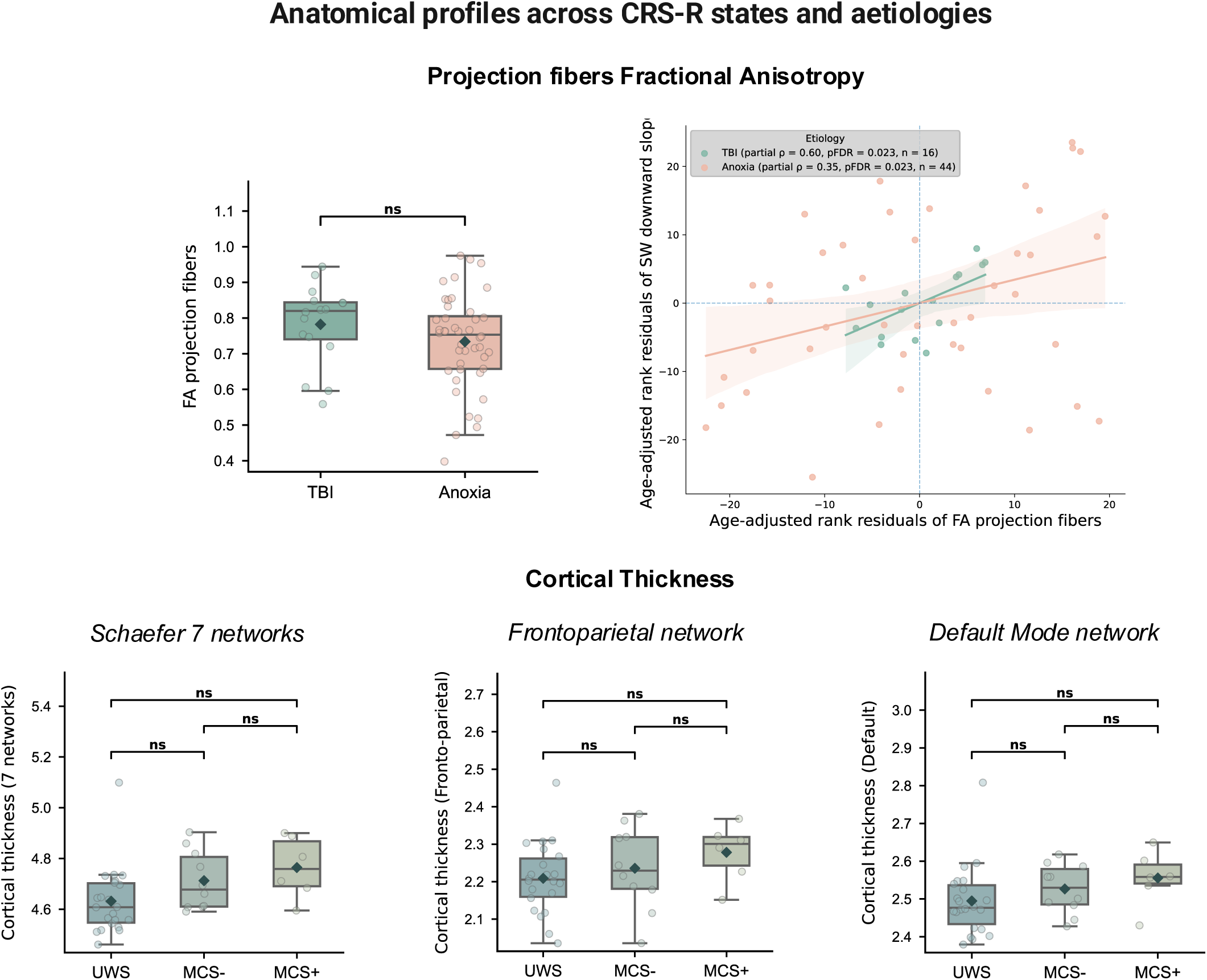
Anatomical profiles across CRS-R states and aetiologies. *Top:* Projection fibers Fractional Anisotropy by etiology group (TBI vs. Anoxia). The scatterplot displays subject-level associations between age-adjusted residuals of FA projection fibers and age-adjusted rank residuals of SW downward slope for TBI and Anoxia patients separately. Regression lines are shown for visualization. Spearman correlation coefficients (ρ) and associated FDR-corrected p-values are displayed for each group. *Bottom:* Cortical thickness by CRS-R states (UWS, MCS- and MCS+) for the Schaefer 7 networks, Frontoparietal network, and Default Mode network. Across the panels, boxplots show the median (central line) and interquartile range (Q1-Q3, box edges), with whiskers extending to the most extreme values within 1.5 × IQR of the box edges. Group-level means are indicated by grey diamonds, while overlaid scatterplots display individual data points (circles). n.s.: non-significant.

**Figure S6:**
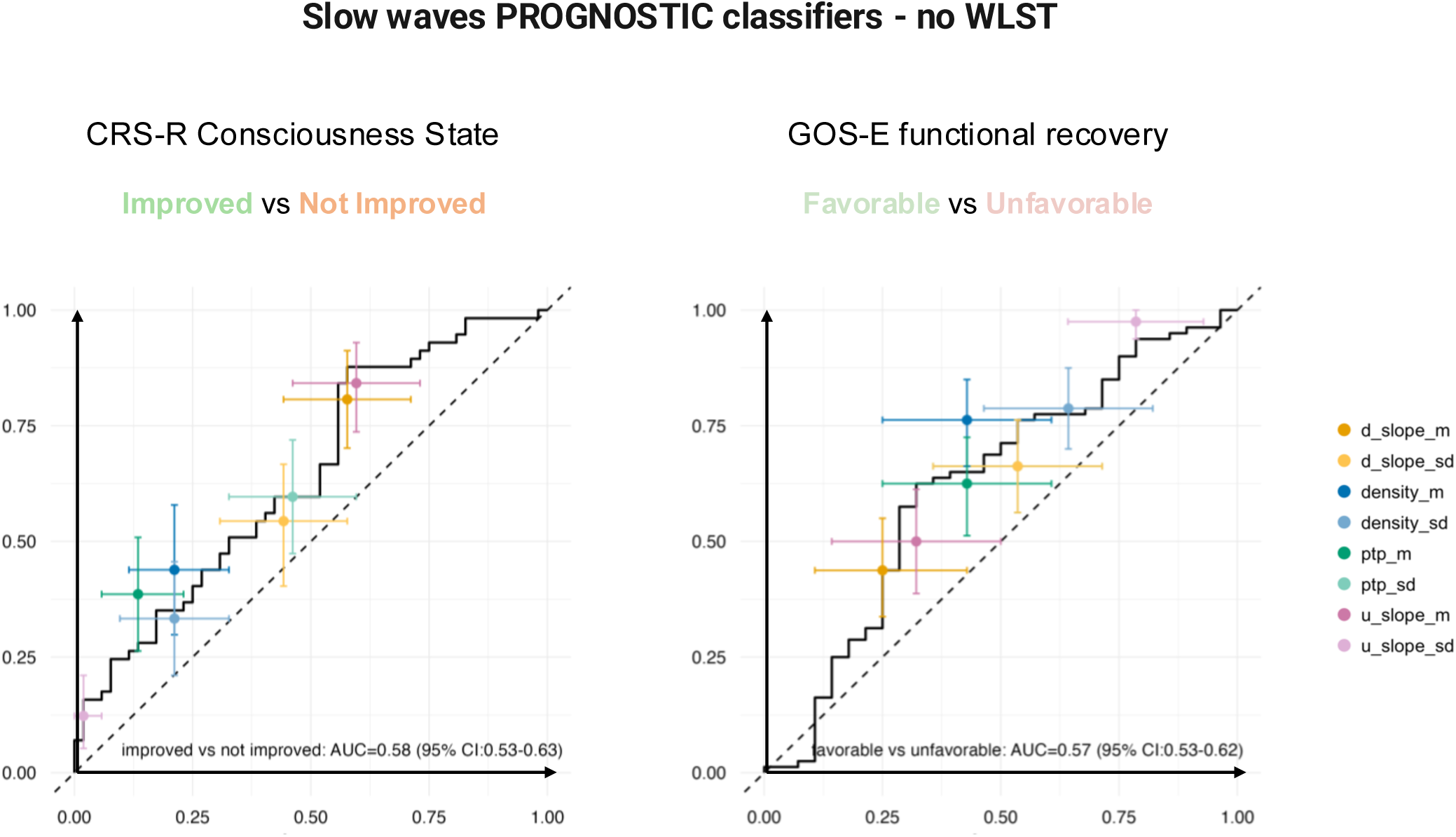
Slow waves PROGNOSTIC classifiers - no WLST. Same as Figure 6 but after removing WLST patients.

